# Spatially-smoothed quantification improves cell typing in imaging mass cytometry datasets

**DOI:** 10.64898/2025.12.16.693927

**Authors:** Reto Gerber, Jake Griner, Daniel Incicau, Silvia Guglietta, Carsten Krieg, Mark D. Robinson

**Author notes:** Corresponding author *Email address:* (Mark D. Robinson).

## Abstract

Accurate cell type annotation in imaging mass cytometry (IMC) and related technologies critically depends on preprocessing steps such as normalization, segmentation, and marker aggregation. More distinct separation between negative and positive signals enables more precise cell boundary inference and more robust marker assignment to single cells. However, inherent limitations in spatial resolution and uncertain cell boundaries can lead to spillover, where signal leaks from one cell to a neighboring cell, distorting marker intensities and leading to incorrect cell type annotation. To address these challenges, we first systematically investigated the impact of spatial resolution and segmentation variability on per-cell marker aggregation using simulated IMC datasets, establishing upper limits for reliable marker separation in cell type annotation. We further analyzed technical biases in large scale studies, demonstrating that appropriate normalization approaches can significantly reduce batch effects without compromising biological variability. Finally, we benchmarked multiple spillover correction strategies across both (semi-)simulated and real IMC datasets. Our results revealed two simple methods: spatial smoothing of intensity images followed by mean marker aggregation, or resampling of cell masks followed by mean marker aggregation and median calculation, both of which improved annotation performance. Other methods often performed worse than the baseline of simple mean aggregation. Together, these findings underscore the central importance of spatial resolution, normalization, and marker aggregation in IMC data preprocessing for accurate single-cell annotation.

## 1. Introduction

Spatial proteomics using multiplexed imaging technologies has revolutionized our understanding of complex tissue architecture by enabling the simultaneous visualization of dozens of protein markers at subcellular resolution. Among these technologies, Imaging Mass Cytometry (IMC) and Multiplexed Ion Beam Imaging (MIBI), have seen wide usage due to their ability to measure around 40 markers simultaneously, which allows fine-grained characterization of cells and tissue, at comparably little cost, enabling interrogation of cellular neighborhoods and interactions [1, 2, 3]. Despite the rich information provided by IMC, extracting meaningful biological insights from these high-dimensional spatial datasets requires processing complex, high-dimensional images, which presents several challenges [4].

A typical computational workflow for IMC data analysis consists of several steps, starting with filtering to reduce technical artifacts such as antibody aggregates or “hot pixels” [4]. Often, simple thresholds are applied to reduce background signal. More sophisticated approaches to improve the signal-to-noise ratio exist, such as IMC-Denoise, which reduces shot noise using differential intensity map-based restoration and self-supervised deep learning [5]. Another commonly applied correction is to reduce signal spillover, caused by isotope impurities and oxidation [6]. After image preprocessing, the next critical and difficult step is cell segmentation. Frequently used tools for cell segmentation in IMC include Illastik [7], Mesmer, [8] and Cellpose [9]. Following segmentation of individual cells, marker intensities are aggregated on a per-cell basis, typically by calculating the mean intensity across all pixels assigned to a cell [4]. This aggregation results in an intensity-per-cell matrix that is used for most downstream analysis. When multiple images are acquired, batch effects can occur and may require additional correction. One approach is to normalize intensities per channel and per image, for example, using z-scores or min-max scaling, or to normalize the aggregated per-cell intensity profiles.

Next, the most common analysis step is cell type annotation, which represents a crucial step transforming high-dimensional marker profiles into biologically interpretable cell identities. Cell type annotation is often performed using either (semi-)manual gating [10] or through unsupervised clustering, followed by manual annotation based on known marker expression patterns [11]. Another increasingly-used set of approaches relies on supervised methods that utilize label prediction based on pretrained classifiers developed from subsets of manually or semi-manually annotated data, enabling more automated and scalable annotation across large datasets [10]. Finally, downstream statistical analysis such as differential abundance or neighborhood analysis can be performed.

Each step in the preprocessing of IMC images contributes to the quality and reliability of IMC data interpretation, but not all of these steps have been equally well studied. IMC image denoising (e.g., IMC-Denoise) and especially segmentation, which is arguably the most important preprocessing step (e.g., Ilastik, Mesmer, Cellpose), have seen large improvements in recent years. The effect of marker aggregation, on the other hand, has not been thoroughly studied, and mean aggregation remains the standard. This is surprising because a well-known issue in cell annotation is lateral spillover, where signal from one cell leaks into a neighboring cell, which cannot be corrected with mean aggregation. Lateral spillover can occur either due to imprecise segmentation, 3D to 2D projections or low spatial resolution and is an inherent issue in IMC and other imaging techniques. Some methods have been developed to improve marker aggregation for multiplexed images, such as REDSEA, which redistributes signal across neighboring cells [12], and Nimbus, a deep learning based marker intensity prediction method pretrained on a large number of semi-manually annotated data [13]. However, both of these methods have not been thoroughly benchmarked and come with some limitations. REDSEA depends on cells touching each other to correct the intensity, which likely works best with expanded cell masks but may show reduced performance on smaller cell masks. Nimbus is a supervised method potentially requiring manually annotated data for fine-tuning to reach decent performance on IMC images.

The paper is organized as follows. Using simulations, we first examine the theoretical properties of mean marker aggregation under different spatial resolutions and segmentation errors. Next, we address batch correction and introduce a new normalization approach for IMC data. We then develop and evaluate two different ways to reduce spillover effects. Finally, we benchmark the performance of the proposed methods alongside other approaches in various scenarios. Developed methods are available at https://github.com/retogerber/cautils.

## 2. Results

### 2.1. Theoretical limit of marker classification depends on image resolution

To investigate the effects of image resolution on mean aggregation of cell markers, we conducted a 3D simulation of small, densely packed, proliferating cells (median area: 31 *µm*^2^; median diameter: 6 *µm*; see Figure 1a, Methods and Supplementary Figure 1). Cells were randomly assigned to one of two classes, either positive or negative for a marker. Additionally, two scenarios of subcellular marker location (“whole_cell” and “membrane”) were simulated. The resulting 3D image stack was subsampled to examine the impact of different spatial resolutions in both the xy and z axes on cell type annotation (i.e., distinguishing positive from negative cells, Supplementary Figures 2 and 3). The area under the receiver operating characteristic curve (AUROC), which gives the probability that a randomly chosen positive cell has higher intensity than a randomly chosen negative cell, was calculated for each slice, and the mean AUROC for each combination is shown in Figure 1. As expected, the AUROC decreases as resolution becomes lower, both for xy and z resolution. Furthermore, if the signal is located only at the cell membrane (Figure 1b), the AUROC is lower than when the signal is found throughout the cell (Figure 1c). These findings suggest that there is a theoretical limit to the quality of cell type annotation in densely packed tissue when using mean aggregation. For instance, for IMC with a xy resolution of 1 *µm* and a z resolution (i.e. tissue depth) of around 4 *µm*, the theoretical limit for this specific simulation is an AUROC of 0.94–0.98, depending on marker location.

**Figure 1.**
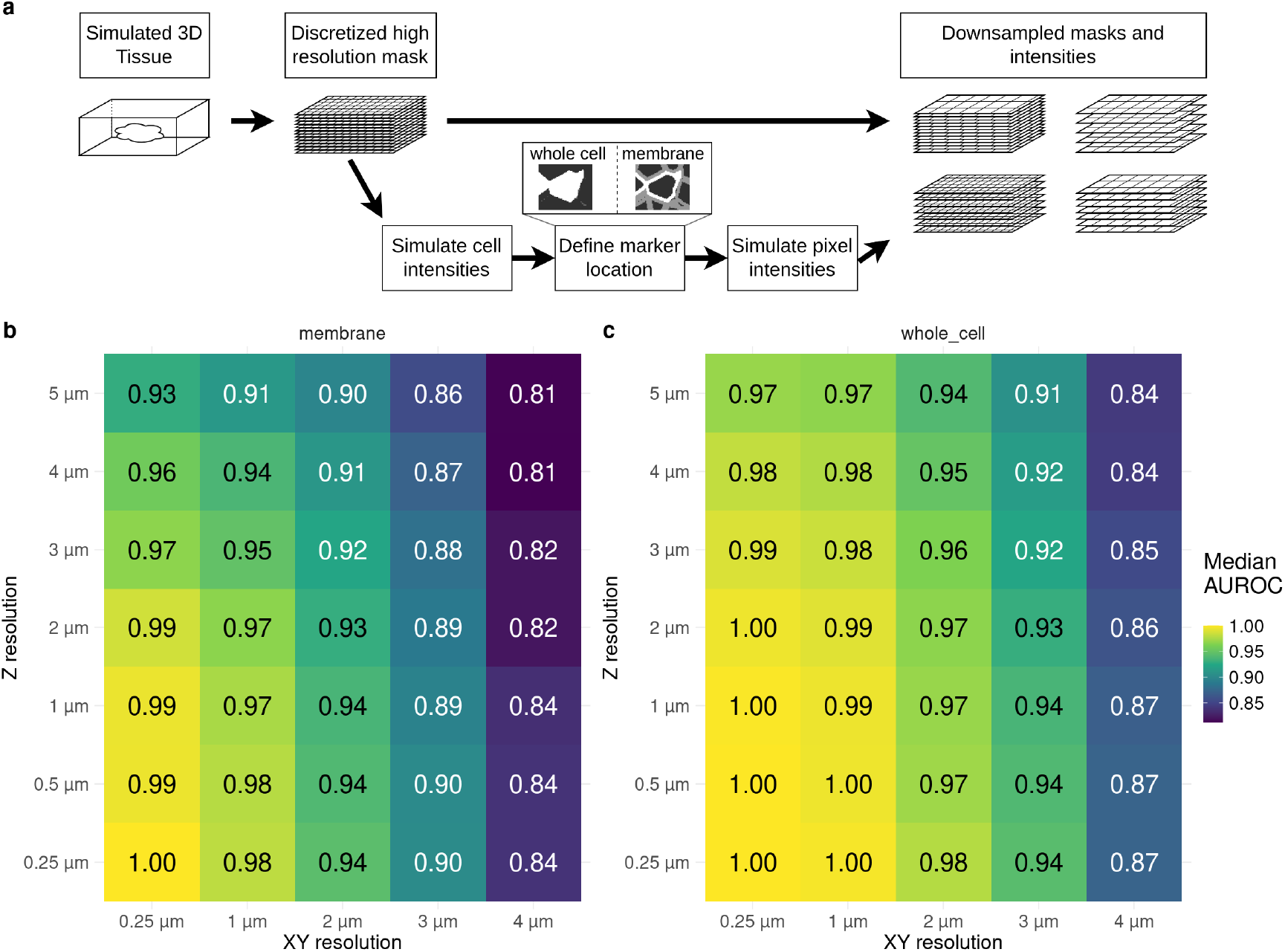
Overview of simulation and results of simulation study. a) Based of a 3D simulation of dense proliferative tissue 2D intensity images and cell masks are generated with two populations, negative and positive, where the signal is either found in the whole cell or only at the membrane. Different spatial resolutions in XY and Z axis are simulated. b-c) Mean AUROC of distinguishing positive from negative cells. The simulation for b) and c) is without noise and with optimal segmentation. b) location of marker intensities at the cell boundary. c) location of marker intensities found in whole cell.

### 2.2. Marker intensity uncertainty depends on cell area and precision of segmentation

In addition to the effects of spatial resolution and 3D structure on marker aggregation, other factors are expected to influence the precision of cell type annotation. For example, cell area is likely to be negatively associated with the stability of mean intensity, which is confirmed by the simulation results. Figure 2a shows how the difference between observed and expected cell intensity depends on cell area. Generally, smaller cells exhibit greater variability, although the average difference does not change across cell areas, indicating that uncertainty in observed intensity is higher for small cells compared to larger ones. Segmentation is another source of error and uncertainty. Figure 2b presents the median AUROC (across multiple slices) for different segmentations and z-resolutions. As shown above, decreasing z-resolution leads to lower performance. When spillover is present, performance decreases compared to the ground truth cell mask, with greater spillover causing a larger drop. If the segmentation mask is too large (“Oversegment”), performance drops slightly, although only boundary cells can be expanded in the densely packed simulation. If the segmentation mask is smaller than optimal (“Undersegment”), performance increases, which can be explained by the fact that in this noise-free simulation, a single center pixel is sufficient for correct cell classification. Additionally, when the segmentation mask is shrunken, smaller cells disappear, further increasing performance (decrease of cell numbers of around 15-20% and 34-38% if shrinking 1 and 2 pixels, respectively; Supplementary Figure 4). These results again provide a theoretical upper limit for marker separability performance under imprecise segmentation.

**Figure 2.**
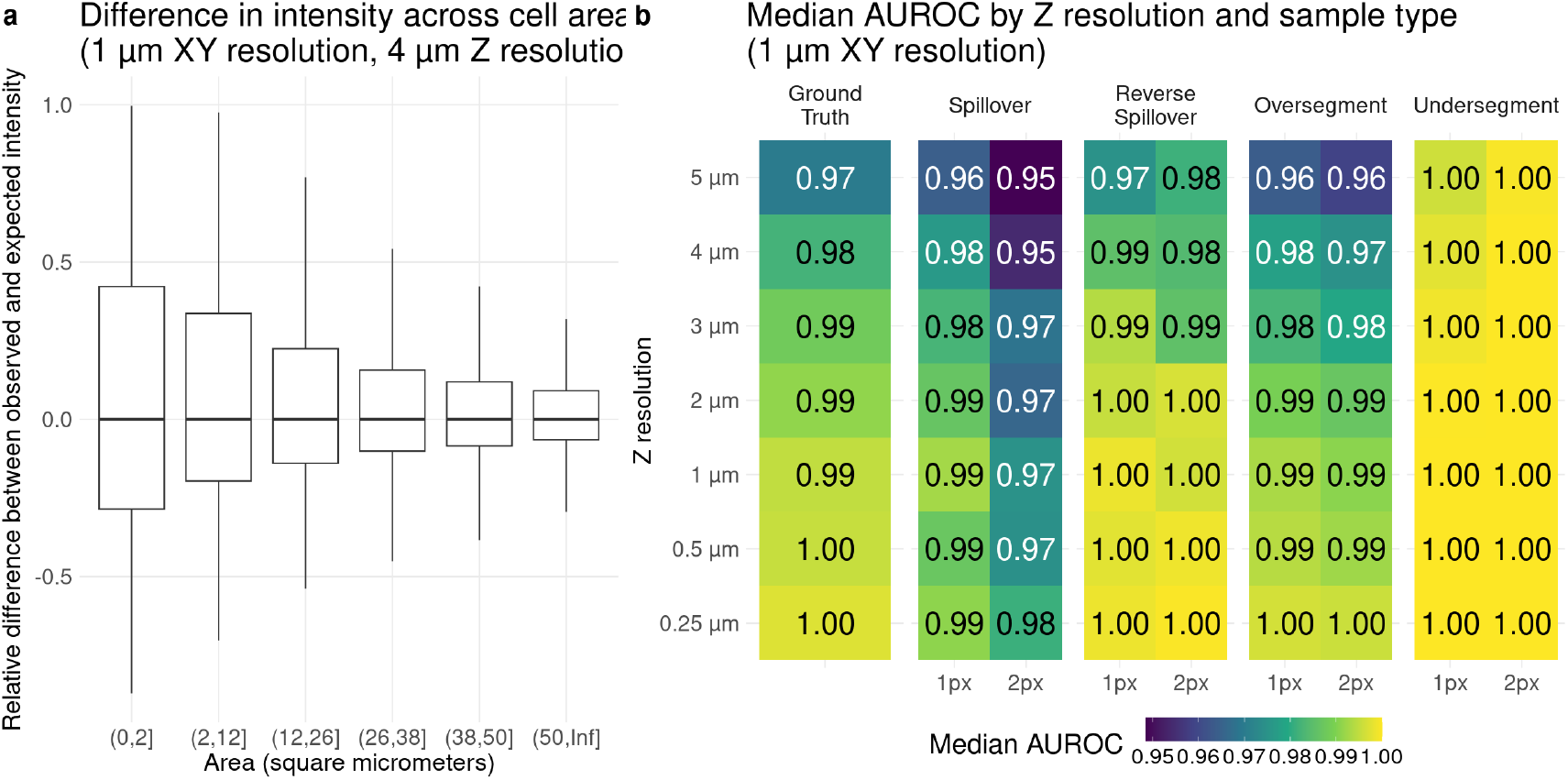
Variables influencing cell annotation. a) shows the relative difference between observed and expected intensity across different cell areas. The number of cells per area bin are similar. b) shows the mean AUROC for different mask errors across different resolutions in z-axis.

### 2.3. Getis-Ord G normalization

Another technical bias commonly observed in larger studies is the presence of batch effects, which are systematic differences arising from non-biological variability caused from unintended technical differences across data acquisitions (for example, fluctuations in antibody staining characteristics over time, or instrument sensitivity). While careful experimental design can reduce batch effects to some extent, completely eliminating them is generally difficult. In image analysis, commonly used normalization methods, i.e., for batch correction at the image level, include min-max scaling (sometimes using quantiles instead of extrema) and z-scores (subtracting the mean and dividing by the standard deviation). However, other approaches exist, such as the local Getis-Ord *G*^*∗*^ statistic, which measures spatial clustering of high or low values and is defined as [14]:

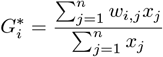

where *x*_*j*_ is the value at location *j, w*_*i,j*_ is the spatial (binary) weight between locations *i* and *j*, and *n* is the total number of locations.

The 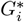 statistic can also be normalized by subtracting the mean and dividing by the standard deviation (see Methods):

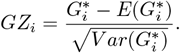

Under the null hypothesis of no spatial signal aggregation, it has been shown to follow a normal distribution with known mean and variance, allowing statistical significance to be calculated. Specifically, the Getis-Ord G statistic is used in hot-spot analysis in geostatistics to identify regions with significantly higher signal than expected by chance. In the context of normalization for image analysis, each pixel can be considered a location, and the Getis-Ord G statistic serves as neighborhood aggregation—a local smoothing—further normalized by the total intensity in the image. In the following, we refer to normalization using the normalized Getis-Ord G statistic as GZ normalization, or simply GZ.

To provide a visual example of GZ normalization, Figure 3a shows an image of CD20 cells with raw, unnormalized intensities, while panel (b) displays the result after GZ normalization. Overall, there is noticeable smoothing, especially in regions of high intensity (the CD20 cells). In the background, intensities appear slightly reduced. The per-cell mean intensities are shown in Figure 3c–d, with ground truth annotations in Figure 3e.

**Figure 3.**
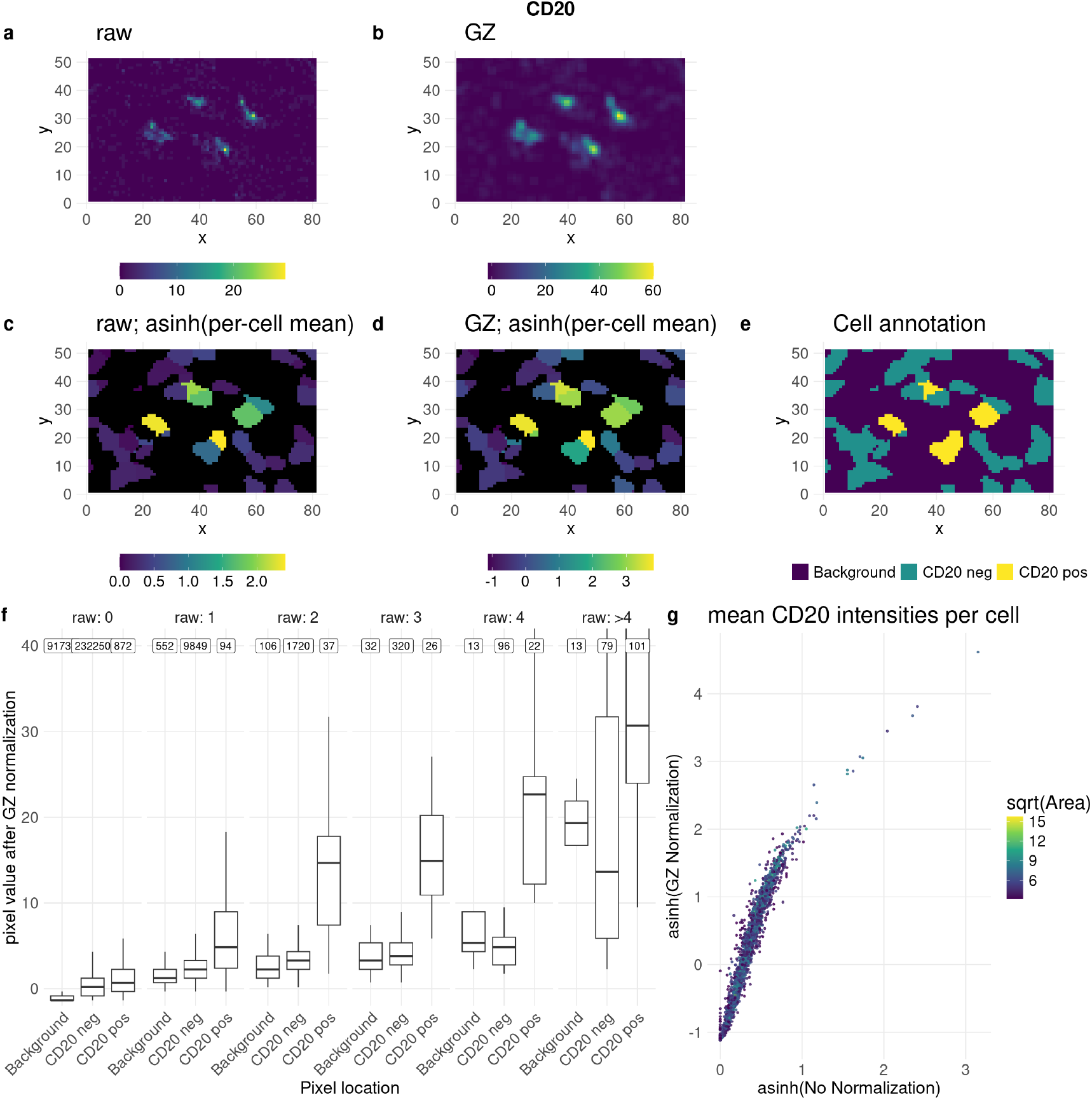
GZ normalization properties for 3D-IMC dataset for CD20 channel. a)-e) example crop depicting some CD20 positive cells. a) raw intensities. b) GZ normalized. c) raw per-cell mean intensities, asinh transformed. d) GZ normalized per-cell mean intensities, asinh transformed. e) ground truth annotation. f) pixel values after GZ normalization, split according to e) and by raw pixel values (columns) according to a). raw: 0 means all pixels that had intensity equal to zero in the raw (unnormalized) image. Numbers at the top show the total number of pixels per bin. g) per-cell raw mean CD20 intensities versus per-cell GZ normalized CD20 intensities.

Quantitatively, low intensity values near high intensity regions are upscaled by GZ normalization (Figure 3f) and remain low elsewhere. After aggregation at the cell level, the unnormalized and GZ normalized values remain quite similar (Figure 3g). In summary, as expected, GZ normalization applies a slight smoothing effect.

### 2.4. Getis-Ord G corrects for batches in IMMUcan dataset

An example of batch effects can be seen in the IMMUcan dataset, which consists of 178 publicly available IMC ROIs across 11 batches [10]. Figure 4a shows low-dimensional embeddings (UMAP) of the entire dataset, split by method. Qualitatively, cells of the same type often appear in regions corresponding to different batches for both ‘mean’ and ‘mean_scaled’, with a seemingly reduced effect for ‘minmaxnorm’, ‘zscore’, and ‘GZ’. Thus, unnormalized mean aggregation does not account for batch effects, and some form of batch correction, such as ‘minmaxnorm’ or ‘GZ’, is needed if the dataset is to be used as a whole. Importantly, the grouping of similar cells (i.e., cells of the same type) is preserved across all normalization approaches (Figure 3b, Supplementary Figure 6). The mean intensities per marker and cell type before and after GZ normalization are very similar (Supplementary Figure 6a), but the variance between batches is, on average, reduced (Supplementary Figure 6b). The Inverse Simpson Indices (ISI), which quantify the local diversity of batches, where higher values indicate better batch integration, are much higher for minmaxnorm, zscore, and GZ normalization than before normalization (Figure 4c, x-axis), indicating reduced batch effects. This result is also observed when stratified by cell type (Supplementary Figure 7), with varying degrees of batch effects per cell type. Additionally, running batch correction using Harmony [15] increases the IS across batches further (Figure 4c, Supplementary Figure 8). Furthermore, the ISI using cell types to quantify diversity (Figure 4c, y-axis) is lower for minmaxnorm, zscore, and GZ normalization than without normalization, indicating better separation between cell types. This is further supported by the Neighborhood Purity (NP) and the Adjusted Mean Shortest Path (AMSP) (Supplementary Table 1).

**Figure 4.**
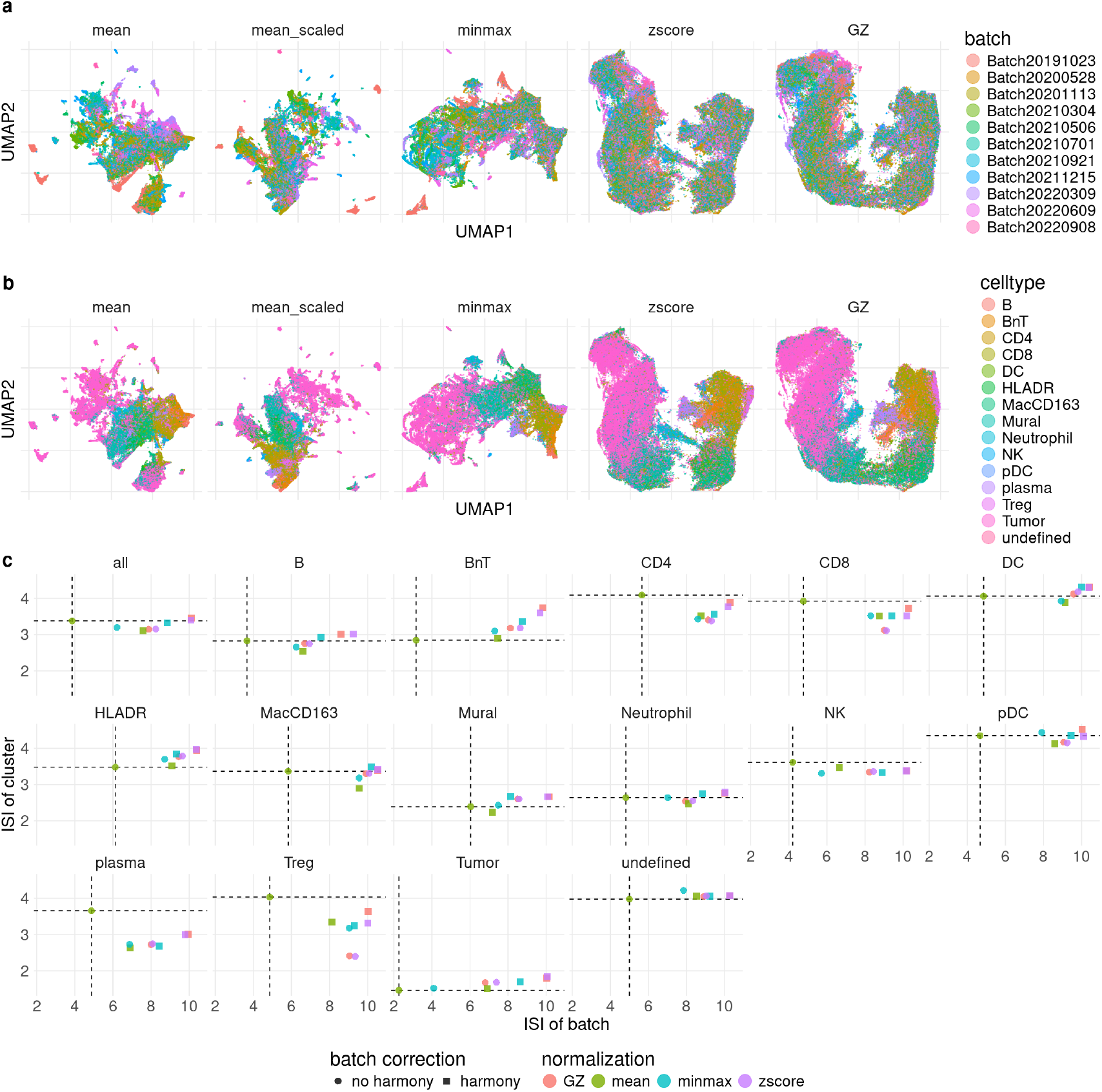
Different normalizations in IMMUcan dataset. a)-b) UMAP for five different normalizations: ‘mean’ (no normalization), ‘mean_scaled’ (minmax scaling at cell level), ‘minmax’ (per ROI min-max normalization), ‘zscore’ (per ROI z-score normalization), ‘GZ’ (per ROI GZ normalization). a) colored by batch. b) colored by celltype. c) Mean Inverse Simpson Index (ISI) per method using either batches (x-axis) or celltype (y-axis) as classes. For the x-axis higher values indicate better batch correction, for the y-axis lower values indicate better celltype separability. The shape represents if additionally batch correction using harmony was performed.

The AUROC for distinguishing a cell type using a single marker is not affected by ‘GZ’ normalization (Supplementary Figure 9), showing that predictive performance remains unchanged. Therefore, using the Getis-Ord G for normalization is a valuable addition to mean marker intensities for batch correction in IMC data.

### 2.5. Spillover correction aggregation

The inherent problem of lateral spillover, resulting from limitations in image resolution (Figure 1) and imprecise segmentation (Figure 2), calls for correction directly at the image level, either before or during marker aggregation. Correcting spillover after mean aggregation is difficult, if not impossible, except perhaps by comparing to neighboring cells.

#### 2.5.1. Proposed methods for spillover correction

We present two simple methods to address some of the issues of lateral spillover. The first method is based on the idea that the local signal within a cell should be consistent, while spillover mainly affects the cell boundary. Starting with an intensity image and its corresponding cell segmentation, the first (optional) step is to apply GZ normalization as described previously (Figure 5b). Next, for each cell border pixel, the potential intensity influence from both outside and inside the cell is calculated (Figure 5c). The difference between these influences is then subtracted from the cell border in the original image (Figure 5d). The resulting corrected intensity image is then subjected to mean aggregation, yielding the final marker value. Two variants for calculating the influence of marker spillover from outside and inside the cell were developed, termed ‘SCI1’ (spillover corrected intensity) and ‘SCI2’ (see Methods). The second method is inspired by cell mask sampling from Bruhns et al. [16], where masks were linearly warped using affine transformations. Multiple resampled segmentation masks are generated (Figure 5e), followed by mean marker aggregation per cell (Figure 5f), resulting in a distribution of possible intensities (Figure 5g). The median of this distribution is then used as the cell intensity. This score is referred to as ‘RATI’ (random affine transformation intensity).

**Figure 5.**
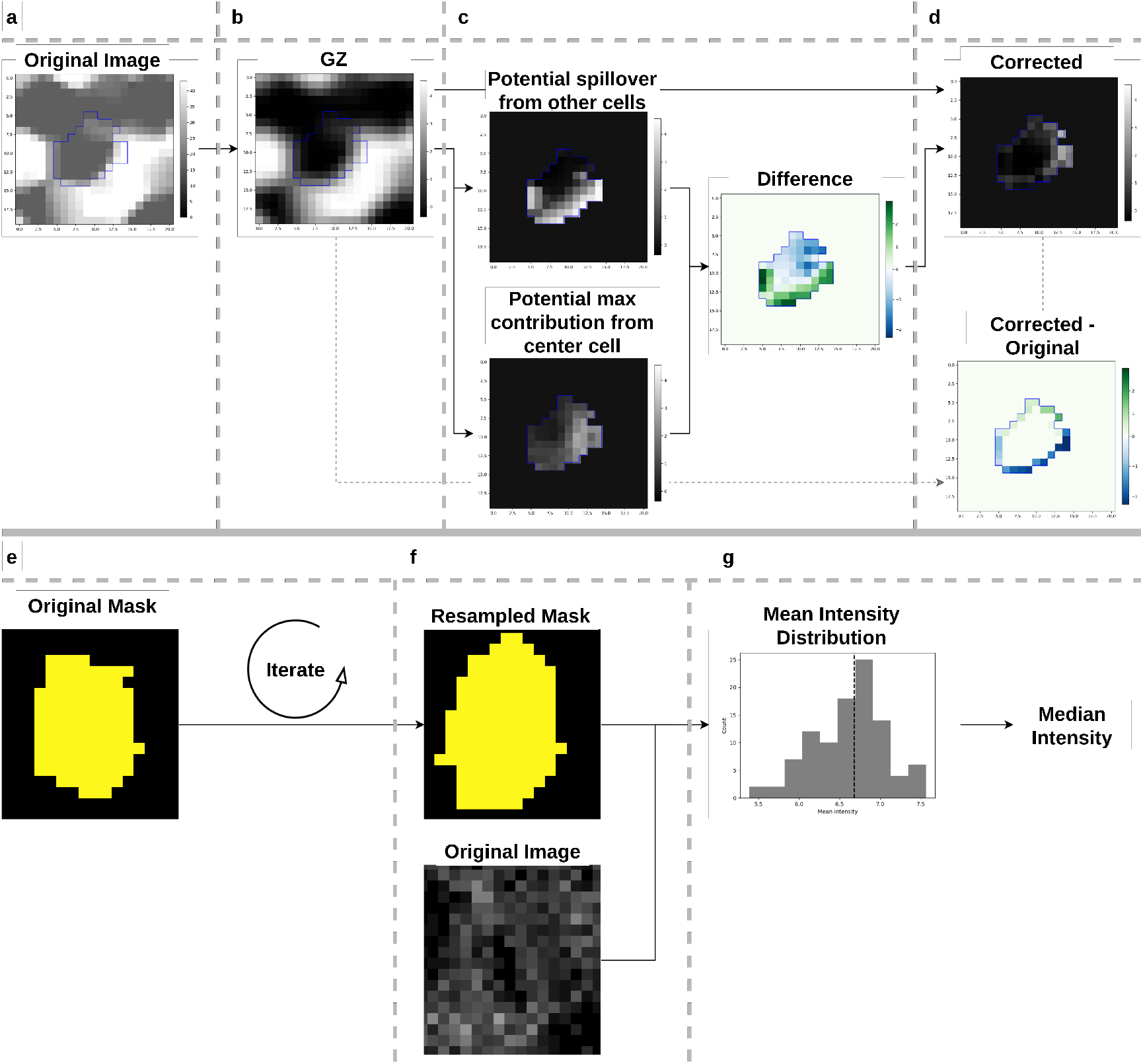
Proposed spillover correction methods, ‘SCI’: a-d, ‘RATI’: e-g. For method ‘SCI’, the original image (a) is first GZ normalized (b), followed by calculation of the potential spillover of neighboring pixels (c, top) and the potential spillover within the cell (c, bottom). The two influences are subtracted and the resulting difference is subtracted from the boundary pixels of the original image (d). For ‘RATI’ the original cell mask (e) is resampled multiple times (f) followed for each iteration by mean aggregation. From the resulting distribution of intensities (g) the median is extracted.

#### 2.5.2. Evaluation of spillover correction

To evaluate the performance of different methods for per-cell and marker aggregation, we conducted a small benchmark using four datasets. For each dataset and marker, the AUROC for distinguishing positive from negative populations, based on ground truth annotations, was calculated and compared to mean aggregation, which serves as the baseline. For the simulated data (Figure 6a, Supplementary Figure 11a), results are shown for two simulated channels and two subcellular marker locations (‘whole cell’ on the left and ‘membrane’ on the right). ‘ch0’ contains substantial noise, while ‘ch5’ contains no noise (see Supplementary Figures 10, 12 and 13). GZ normalization alone performs slightly worse in terms of AUROC, but the proposed scores slightly improve performance when the subcellular location is ‘whole cell’. For the ‘membrane’ subcellular location, SCI2 shows increased performance over all other methods. RATI also outperforms mean aggregation and achieves similar performance to either SCI1 or SCI2. Interestingly, for many markers in the 3DIMC, IMCIF, and Lunaphore datasets (Figure 6b–d, Supplementary Figures 14-21), GZ normalization alone increases performance in nearly all cases. Since GZ normalization essentially performs local smoothing and scaling, the ‘blur’ method, which is simply image convolution with a box filter, is also shown and yields exactly the same AUROC as GZ normalization. SCI1 and SCI2 show some improvement in certain cases, while a decreased AUROC is observed for markers located at the cell membrane (e.g., CK19) or at boundaries between spatial domains (e.g., SMA). For “easy” cases, with AUROC values close to one, the proposed scores remain very high as expected (e.g., channel “E-Cad” for the IMCIF dataset in Figure 6c). RATI performs at least as well as mean aggregation across all conditions. The neighborhood for GZ normalization can be defined in various ways, depending on the distance at which pixels are considered neighbors. In Figure 6, GZ normalization uses all pixels with a Euclidean distance less than 1.5 as neighbors, corresponding to the eight nearest neighbors. As the neighborhood distance increases, performance varies depending on the marker (Supplementary Figure 25). For example, in the Lunaphore dataset, CD3 performance initially improves with increasing distance, then declines. The optimal distance appears to be highly marker-dependent, with some cases not benefiting from smoothing. Additionally, tissue density and the expected neighboring cell types play a role. In simulations with very dense tissue and random cell type labeling, any smoothing reduces performance (Supplementary Figure 25a). Conversely, in the less dense tissue of the 3D IMC dataset (at least in terms of segmentation mask), smoothing improves performance for all markers (Supplementary Figure 25b). There is also a dependence on the cell segmentation mask, since masks that are smaller than optimal generally decrease performance (Supplementary Figure 26).

**Figure 6.**
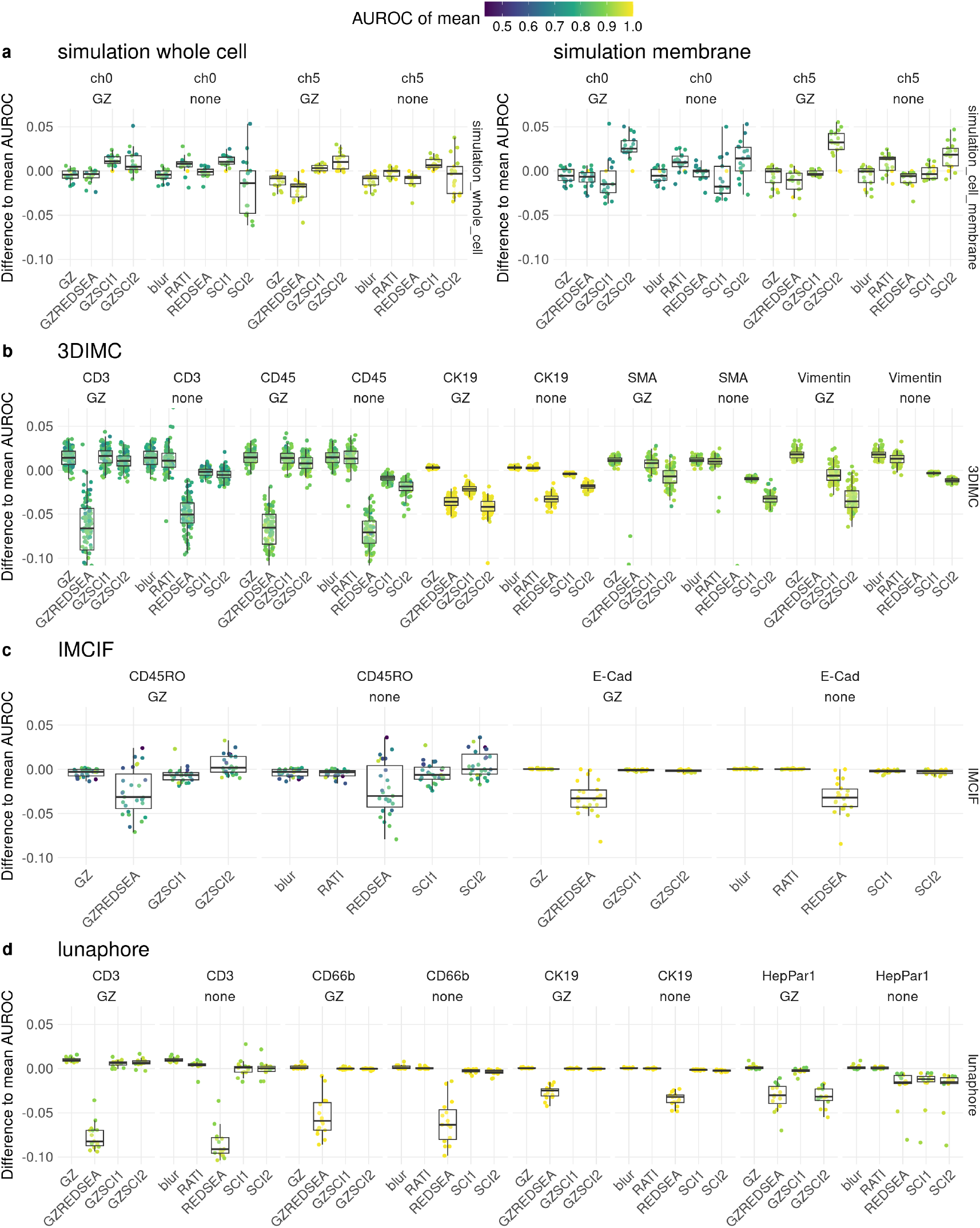
AUROC difference of methods to ‘mean’ baseline for different methods, normalization, marker and data set. Included methods have been tested with GZ normalization (‘GZ’ prefix) or without. a) simulation of high noise channel (‘ch0’) and no noise channel (‘ch5’) for marker location whole cell (left) and boundary only (right). b) 3D IMC dataset for 5 different channels with different subcellular localization. c) IMC IF dataset for two matching channels.

Other methods evaluated include REDSEA and Nimbus. The REDSEA method, both unnormalized and with GZ normalization, consistently performs worse across all datasets and markers. The exact reason for this requires further investigation, but it appears that REDSEA overcorrects the marker intensities. Even poorer performance is observed with Nimbus when using the pretrained model weights (Supplementary Figures 12 - 20). Fine-tuning would likely improve Nimbus’s performance, but this was not done in the present study.

Examples of how AUROC scores correspond to marker distribution and other performance metrics are shown in Supplementary Figures 29 - 31. In short, even a modest increase in AUROC can be a substantial increase in classification performance in terms of F1-score.

To understand why GZ normalization improves performance, we calculated the distances between positive and negative populations (i.e., average intensities). These distances show a slight increase with GZ normalization compared to no normalization (Supplementary Figure 23a), which corresponds to higher AUROC scores. Additionally, when spillover is simulated, these distances remain very similar for both unnormalized and GZ normalized data (Supplementary Figure 23b).

Since the RATI method depends on multiple parameters, a small parameter sweep was done to investigate the effects of the number of iterations and the sampled parameters in the affine transformation (translation, rotation, scale, shear). In general, even with just 10 iterations, an increased AUROC is visible in many cases with some additional increase with more iterations for some situations (Supplementary Figure 27), and surprisingly, translation seems to have the most influence on improving results (Supplementary Figure 28).

To see how the performance changes if the random spillover is simulated by slightly changing segmentation masks (see Methods, Figure 2 and Supplementary Figure 4), the obtained AUROC scores were compared between correct segmentation and sampled segmentation, which can be interpreted as a stability of the score under misssegmentation (Supplementary Figure 24). However, no systematic difference between GZ normalization and no normalization were detected.

## 3. Discussion

Cell type annotation in IMC relies heavily on cell segmentation and marker aggregation. Segmentation is challenging and prone to errors, which are then propagated to the next step: marker aggregation. A common issue is lateral spillover, where the signal from one cell leaks into a neighboring cell. Additionally, batch effects in large studies can complicate cell type annotation and interpretation.

Interestingly, applying GZ normalization not only reduced batch effects but also improved cell type annotation performance. Closer inspection suggests this effect is due to local image smoothing. The optimal degree of smoothing depends on the specific marker, but generally, using the queen neighborhood (i.e., eight nearest neighbors) resulted in consistent improvements across various datasets and markers. This observation aligns with previous findings, such as those from PIXIE [11], where a Gaussian blur enhanced annotation performance. The batch correction capability, on the other hand, appears to stem from the z-score normalization term in GZ normalization, as simple z-score normalization has a similar effect on batch correction in the IMMUcan dataset.

The observation that increasing the amount of blurring to a seemingly excessive degree (e.g., a radius of 5 *µm*, which is the size of some cells) may be explained by the spatial distribution of the marker in the tissue. For example, in the 3D IMC dataset, SMA is often found only on one side of many cells, and smoothing likely increases the signal across the entire cell. Conversely, if the spatial signal distribution is random, as in the simulation, any smoothing reduces the AUROC.

The developed scores, SCI, were designed to reduce spillover effects and tuned using simulated data, did not lead to improvement, suggesting that further investigation is needed. Since the simulation used optimal segmentation, an obvious next step is to examine the role of segmentation and possibly develop methods that better account for segmentation mask uncertainty. Additionally, simulations with less random localization of cell types could help clarify the effect of neighboring cells of similar type on annotation.

The score based on mask resampling (RATI), however, did improve performance. Although random sampling followed by two rounds of aggregation can be considered a form of smoothing, it may not be linear. The parameter space for affine transformations and the number of samples were not thoroughly explored, indicating room for improvement by optimizing these parameters, either globally or on a case-by-case basis. The randomness of this score is also not ideal, but setting random seeds, increasing the number of samples, or systematically determining transformations (perhaps with a parameter grid) could mitigate this issue. Additionally, this bootstrapping approach could provide diagnostics for cell missegmentation (e.g., standard deviation of intensities).

One major difficulty in evaluating cell type annotation methods in IMC is the limited number of available data sets where the ground truth has been established using orthogonal methods (e.g., IF) to reduce biases. While simulated data with known ground truth was used, it does not capture all the complexities of real IMC data. Similarily for the semi-simulated data which was generated based on observed IF images.

## 4. Methods

### 4.1. Simulation

An input mesh file was manually created in Blender [17] consiting of a set of regularly spaced spheres with a volume comparable to cells. simucell3d was then used to simulate dynamically growing cells in 3D [18]. After running the simulation the resulting mesh is a densely packed clump of cells. The resulting mesh was then transformed to a 3D mask using pyvista [19] by thinly slicing the mesh and coloring by cell id. Additionally a second mask was created which only included the boundary of a cell by subtracting the eroded (with a 3×3 kernel) mask from the original mask. Image processing was done in Python using opencv [20]. Downsampling of masks was done by nearest neighbor interpolation for the x-y axis and maximum occurrence for the z-axis (projection). Cells were defined to be either positive or negative for a marker by a random draw from a bernoulli distribution with probability of success equal to 0.5. The mean expression of the cell types (positive or negative) was then defined to come from a random draw from a truncated normal distribution (followed by an exponential transformation) with parameters only differing for the mean in the two groups (for ‘ch0’: mean of negative population=0.5, mean of positive population=1.3, standard deviation=0.3. for ‘ch5’: no randomness, mean expression of negative population=0, mean expression of positive population=100). The actual counts image was then constructed in two steps. First, the mean intensity for each pixel was created based on present cell and its corresponding random draw of the mean expression, with additionally added offset and simplex noise [21]. Next, depending on the needed resolution, the mean intensity image was downsampled. Second, the actual counts for each pixel was simulated by a random draw from a poisson distribution with the size parameter corresponding to the mean intensity specified in the previous step. For the evaluation of the theoretical properties the second step was omitted and an unsampled (no offset and no simplex noise) mean intensity image was used.

### 4.2. 3D IMC processing

The 3D IMC datasets ‘MainHer2BreastCancerModel’ and ‘SecondHer2BreastCancerModel’ were downloaded from https://zenodo.org/records/5782846. Segmentation was done using cellpose v4 [22] using for the nuclear channel the mean of markers Ir191, Ir193 and Histone H3, and for the cell membrane channel the mean of markers E/P Cadherin and panCK. The ground truth marker annotation (i.e. if a cell is positive or negative for a marker) first the mean intensity from the 3D image stack was calculated followed by asinh transformation with a cofactor of 50 and z-score normalization. Then a semi-manual gating strategy was employed by visually inspecting the thresholding results of multiple automatic thresholding methods and choosing the most appropriate one.

### 4.3. IMC - IF processing

The IMC-IF dataset was downloaded from https://zenodo.org/records/7576005 and the original scripts from the publications were run to obtain a single integrated image for the IMC registered to the IF. The IF image was processed in the following way: median filter with a kernelsize of 5, tophat convolution with a round kernel of diameter 31, setting background signal - defined via thresholding - to zero. Segmentation was done using cellpose v4 two times, once for nuclear only (DAPI channel) and once for whole cell segmentation (DAPI and E-Cadherin channels). The IMC images were segmented cellpose v4 again for nuclear only (DNA channel) and whole cell (DNA and E-Cad channels). Nuclear and whole cell segmentations were matched by considering the only objects that are shared nearest neighbors with a maximum distance of 5 *µm* between centroids and where the nuclei mask is inside the whole cell mask (with a tolerance of 4 square *µm*). Across modalities the matching was purely done based on shared nearest neighbors of the nuclei with a maximum distance between centroids of 2 *µm*. Classification of IF markers (CD45 and E-Cadherin) into positive and negative was done by first extracting features using Nyxus [23], manual annotation of a set of cells using napari-feature-classifier [24] followed by training a random forest classifier and prediction of all remaining cells.

### 4.4. Lunaphore COMET processing

Two slides of formalin fixed, parrafin embedded liver tissue were deparaffinized and subjected to heat induced epitope retrieval before being imaged with the COMET platform from Lunaphore. A total of 20 antibodies targeting the following were used: HepPar1, CK19, CD66b, CD32b, HLA-DR, CD3,CD8, CD14, CD20, CD68, CD31, GzmB, aSMA, CD16,CK, Vimentin, E-Cadherin, CD11b, CD4, CD45 and DAPI. Autofluorescence is measured in both the TRITC and Cy5 channel for subtraction from subsequent images. Final images were then acquired across 18 rounds of cyclic immunofluoresence, including negative controls. These data were acquired through the Lunaphore COMET Access Lab.

The autofluorescent subtracted images provided by lunaphore were used as the starting point. Cell segmentation was done using cellpose. Regions containing image artifacts (e.g., air bubbles, folds, etc.) were manually selected and excluded from downstream analysis. IMC data was acquired from adjacent tissue slices containing the same antibodies as for fluorescent images. The semi simulated IMC images were then created by first downsampling the COMET images to IMC resolution (1 micron). This was followed by matching the histograms of channels between IMC and COMET. Next, a value of 0.5 was added to all pixels, followed by drawing a random sample for each pixel from a poisson distribution with the expected value equal to the previous intensity value. Classification into positive and negative for a marker was done by first calculating features using Nyxus followed by manual annotation using napari-feature-classifier and training of a random forest classifier, and finally prediction.

### 4.5. IMMUcan processing

The IMMUcan dataset was downloaded from https://zenodo.org/records/12912567. The provided cell masks and cell type annotation were used. GZ normalization was done as previously described. Scaling per batch was done by cropping all values between the 1 and 99 percentiles and scaling between 0 and 1. Cell type marker extraction was done on the unnormalized mean asinh transformed intensities using scran [25]. Batch correction was performed using Harmony [15]. AUROC calculation was done using pROC [26] with the reference being a specific cell type and the score the associated single marker value, either the unnormalized or the GZ normalized variant. The Inverse Simpson Index (ISI), Neighborhood Purity (NP) and the Adjusted Mean Shortest Path (AMSP) were calculated using the R package poem [27].

### 4.6. Mask sampling

To simulate spillover the mask of individual cells was slightly changed based on the following procedure: First a cell is randomly selected, next an operation - dilation or erosion - is randomly selected, then a quadrant (top left, top right, bottom left, bottom right) is randomly selected, then a range (1 or 2 pixels) is randomly selected. The specified operation is then applied to the single cell by essentially removing (erosion) or increasing (dilation) the cell mask according to the sampled parameters. Then for evaluation only the specifically modified cells are considered. For the undersegmentation each cell was individually is eroded by either 1 or 2 pixels. For the oversegmentation each cell was individually dilated by either 1 or 2 pixels, but only if the space is not yet occupied by other cells, meaning for the densely packed simulated tissue this was mostly the case at the border of the image.

### 4.7. Spillover correction scores

The proposed spillover correction broadly works as shown in Figure 5. After GZ normalization as described above, this is followed by estimation of intensity image “A” which is the potential spillover from other cells and estimation of intensity image “B” which is the potential max contribution from the center cell. This is followed by calculating the mean between the two and subtracting it from the intensity. Two different versions of calculating “A” and “B” have been developed, “SCI1” and “SCI2”. For both versions, first two separate images and mask are created: For “A”, the mask (“AM”) is equal to one for all pixels that are not the center cell and zero otherwise, the intensity image (“AI”) is set to zero for all pixels belonging to the center cell and otherwise unchanged. For “B”, the mask (“BM”) is equal to one for pixels that are the center cell and zero otherwise, the intensity image (“BI”) is zero for non-center cell pixels and otherwise unchanged. Then “SCI1” and “SCI2” process “A” and “B” differently. For “SCI1”, “AM” and “AI” are both filtered with a gaussian filter with size 5, followed by dividing “AI” by “AM” (for weighting) which results in “A”. Similarly for “B”, which is obtained by gaussian filtering (kernel size: 9) “BM” and “BI” followed by dividing “BI” by “BM”. For “SCI2” calculating “A” and “B” is a bit more involved. First, a set of 8 Gabor like kernels are created with the following parameters: size=7, sigma=3, theta=0 to 2*pi (8 evenly spaced), lambd=10, gamma=5, psi=pi/2 (opencv function getGaborKernel). All gabor kernels are then applied to “AM”,”AI” and “BI” resulting in images stacks “AMG”,”AIG” and “BIG”, each with 8 channels representing the individual filters. Next, for each pixel in “AMG” the maximum value across channels is evaluated, and if this is ambiguous, the minimum is evaluated and the channel that is perpendicular is used in terms of the theta parameter of the gabor filters (i.e., if minimal for kernel with theta=0 the kernel with theta=pi is used). Then, the “BIG” values corresponding the just evaluated max “BMG” are used resulting in “B”. This essentially results in taking the values perpendicular to the segmentation mask. For “AIG” the maximum intensity value across channels is used, giving “A”, the maximum local contribution possible inside the cell. The resulting processed “A” and “B” images are subtracted in the same way for both “SCI1” and “SCI2”.

For the random affine transform intensity (‘RATI’), the complete cell mask is resampled by randomly iterating over cells, and for each cell applying an affine transformation with randomly chosen parameters. This is followed by a closing operation to remove gaps in fully filled regions. The parameters for the affine transform are sampled individually from an uniform distribution with the following limits: scale from 0.8 to 1.2, translation (in pixels) from -1 to 1, shear from -0.1 to 0.1, rotation (in degrees) from -15 to 15.

### 4.8. Evaluation

The Area Under the Receiver Operator Curve (AUROC) is a performance metric for binary classification problems. It measures the ability of a classifier to distinguish between positive and negative classes. The AUROC is defined as the probability that a randomly chosen positive instance is ranked higher than a randomly chosen negative instance by the classifier.

Mathematically, the AUROC can be expressed for a set of *n*_+_ positive and *n*_−_ negative samples with scores *s*_*i*_ as:

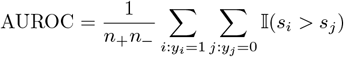

where 𝕀 is the indicator function, *y*_*i*_ is the true label, and *s*_*i*_ is the predicted score. The F1 score is defined as follows:

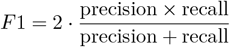

where:

- Precision 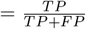
- Recall 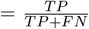

with TP = true positives, FP = false positives, FN = false negatives. The threshold for classification is chosen to maximize the F1 score.

### 4.9. Getis-Ord G

The local normalized Getis-Ord *G*^*∗*^ statistic is defined as [14]

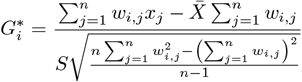

where:

- *x*_*j*_ is the value at location *j*,
- *w*_*i,j*_ is the spatial weight between locations *i* and *j*,
- *n* is the total number of locations.
- 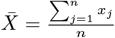
- 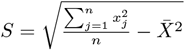

The neighborhood of a location is defined as its “Queen” neighborhood, i.e. for a regular 2D grid of locations its 8 neighbors. The described GZ normalization is simply calculating this statistic for each pixel.

## 5. Data availability

The IMC-IF dataset was downloaded from https://zenodo.org/records/7576005. The IMMUcan dataset was downloaded from https://zenodo.org/records/12912567. The 3D IMC datasets ‘MainHer2BreastCancerModel’ and ‘SecondHer2BreastCancerModel’ were downloaded from https://zenodo.org/records/5782846. The lunaphore dataset can be downloaded from: https://www.ebi.ac.uk/biostudies/bioimages/studies/S-BIAD2494.

## 6. Code availability

The analysis scripts are available at https://github.com/retogerber/celltyping_imc (and on zenodo: https://doi.org/10.5281/zenodo.17930566). Code to calculate GZ normalization and RATI can be found at https://github.com/retogerber/cautils.

## 7. Author contribution

Data acquisition: JG, CK. Data analysis: RG and MDR. Conceptualization and data interpretation: RG and MDR. Funding: MDR. Writing: SG, CK, MDR, RG, JG, DI.

