## Supplementary Information for "Spatially-smoothed quantification improves cell typing in imaging mass cytometry datasets"

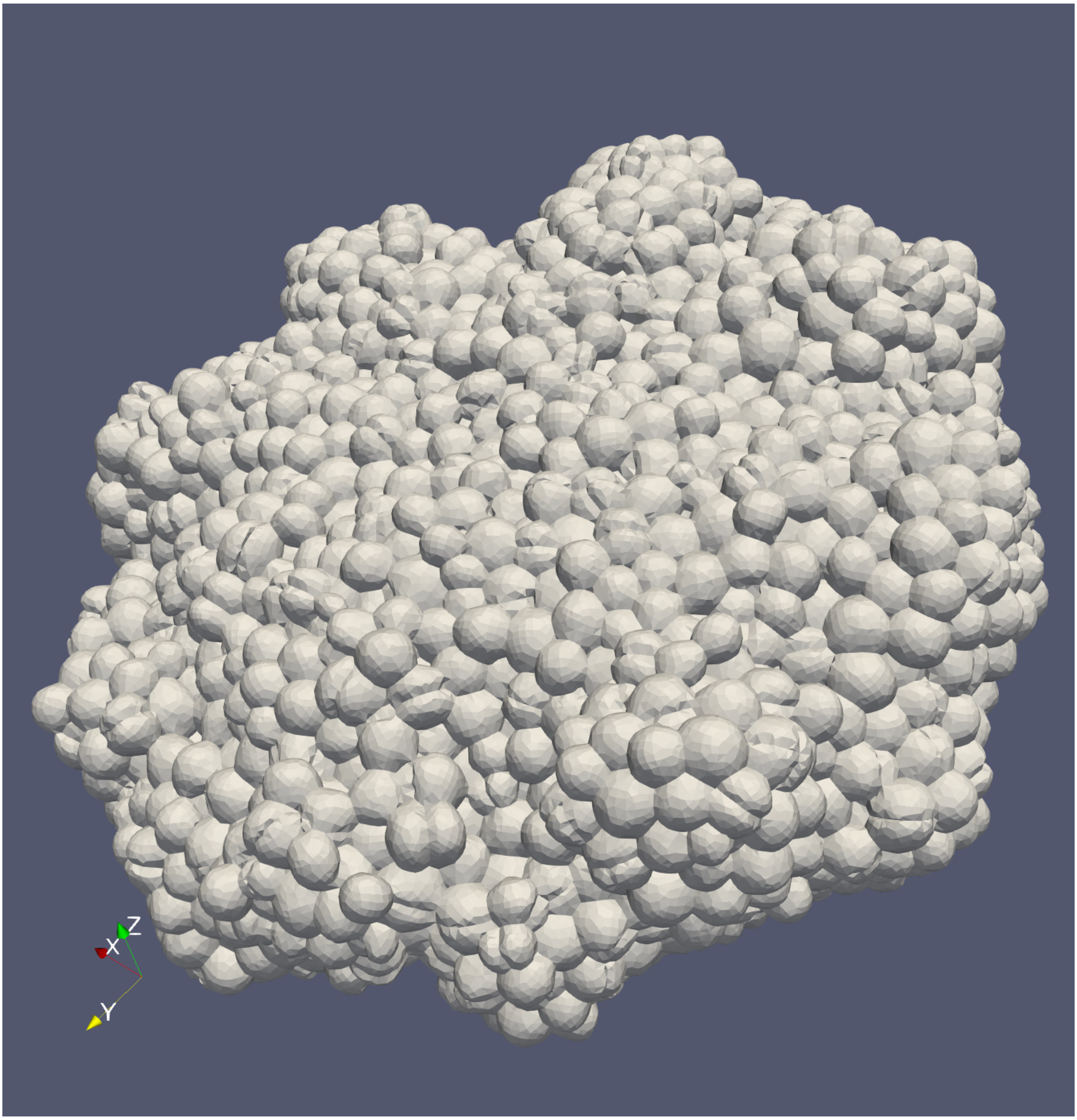


Supplementary Figure 1: 3D rendering of simulation

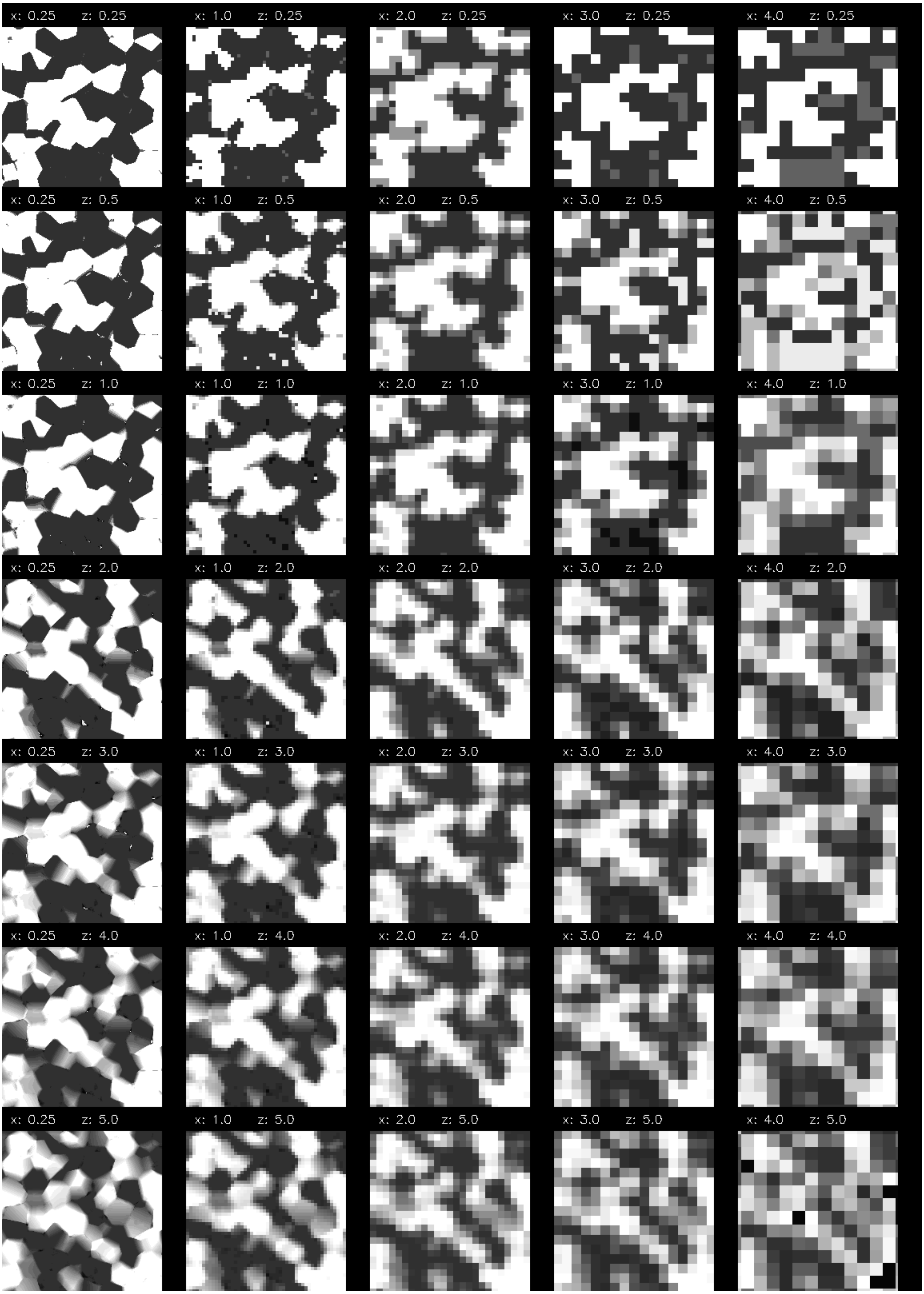


Supplementary Figure 2: Example image crop of the simulation showing the intensity of a noise-free marker across a set of spatial resolutions in xy and z-axis.

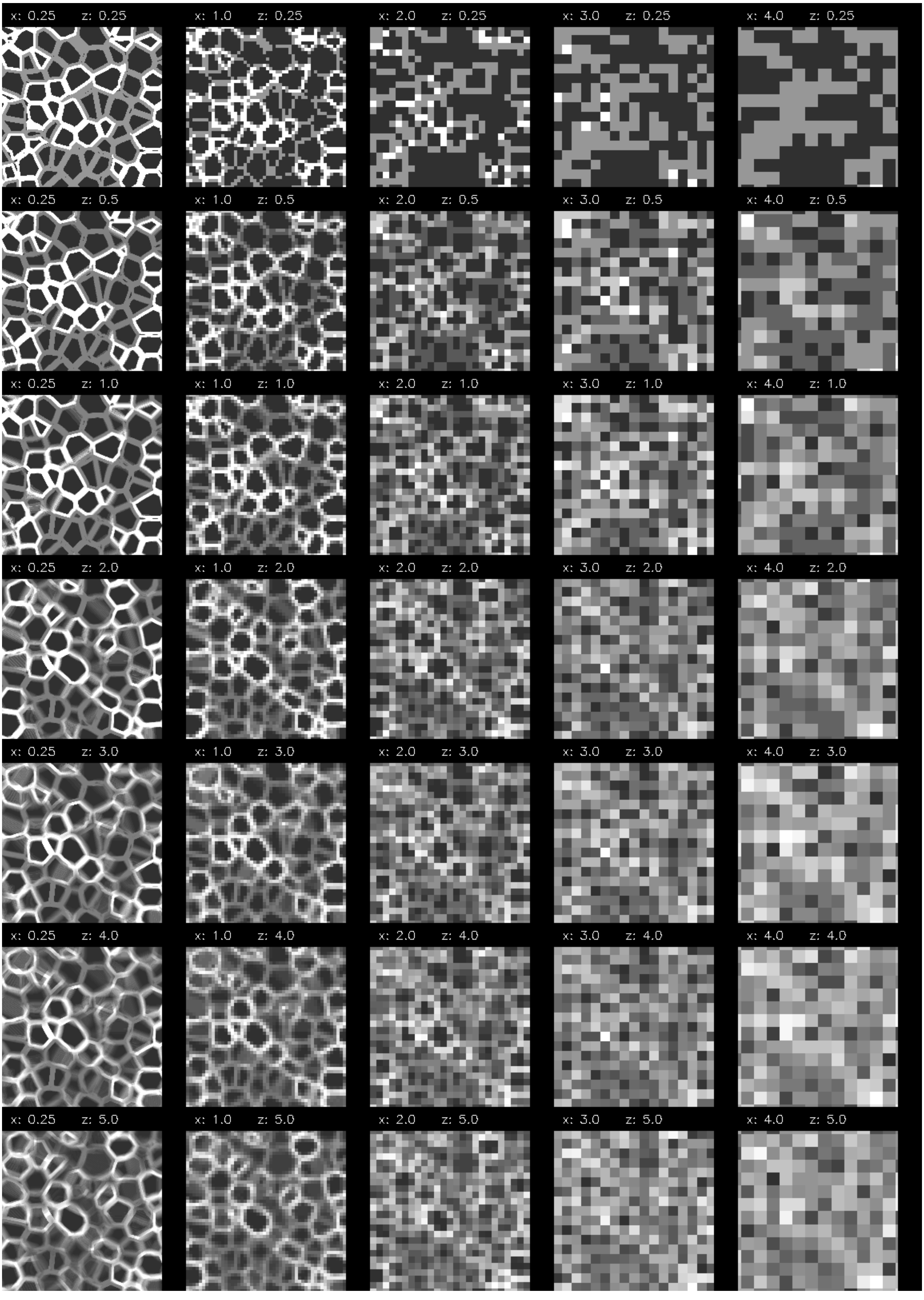


Supplementary Figure 3: Example image crop of the simulation showing the intensity of a noise-free marker across a set of spatial resolutions in xy and z-axis.

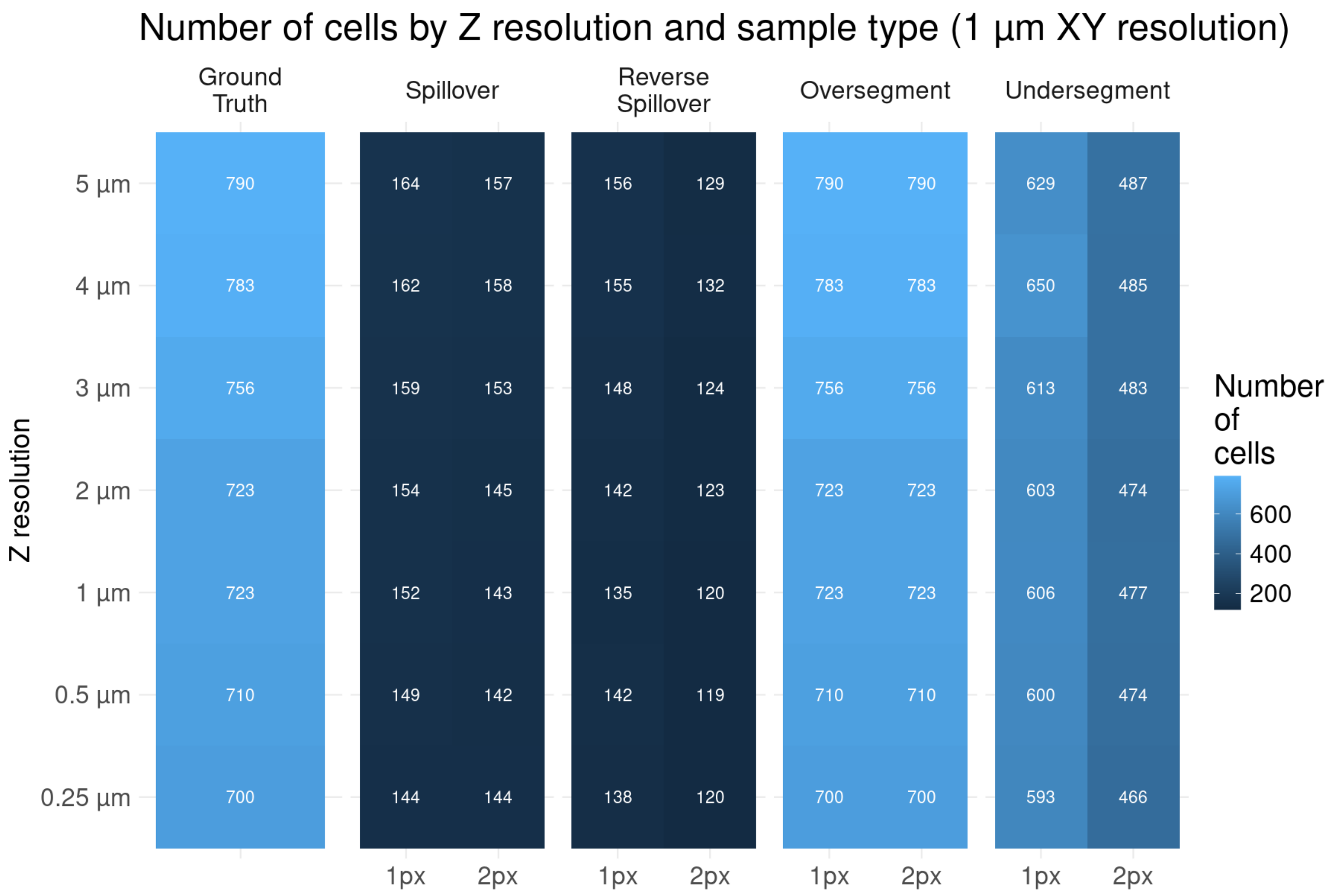


Supplementary Figure 4: "Mean number of cells in the simulation for different mask errors across different resolutions in z-axis and fixed xz-resolution (1 micron)"

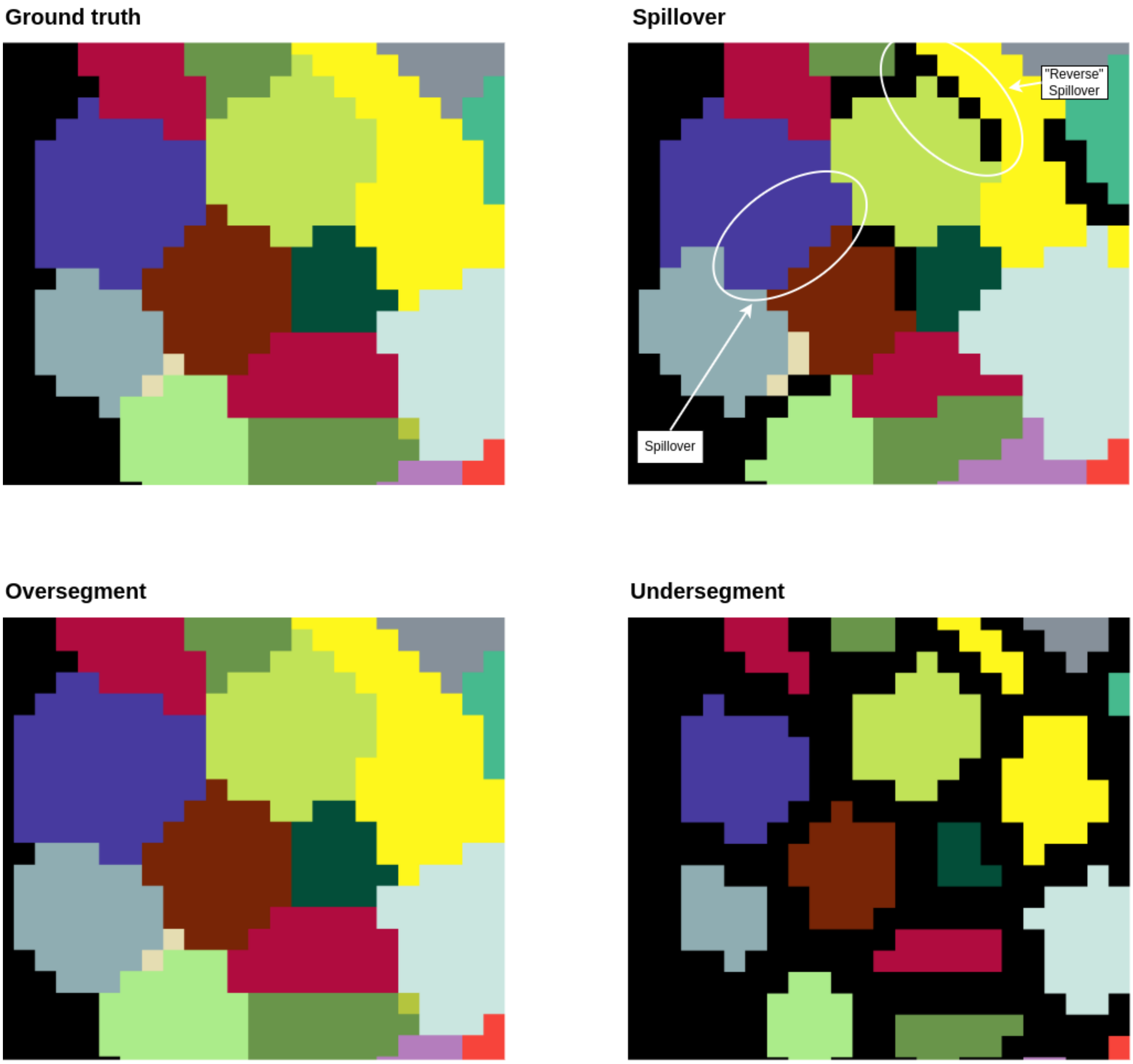


Supplementary Figure 5: Example of applied mask errors. Top left: ground truth. Top right: Spillover. Bottom left: Oversegment. Bottom right: Undersegment.

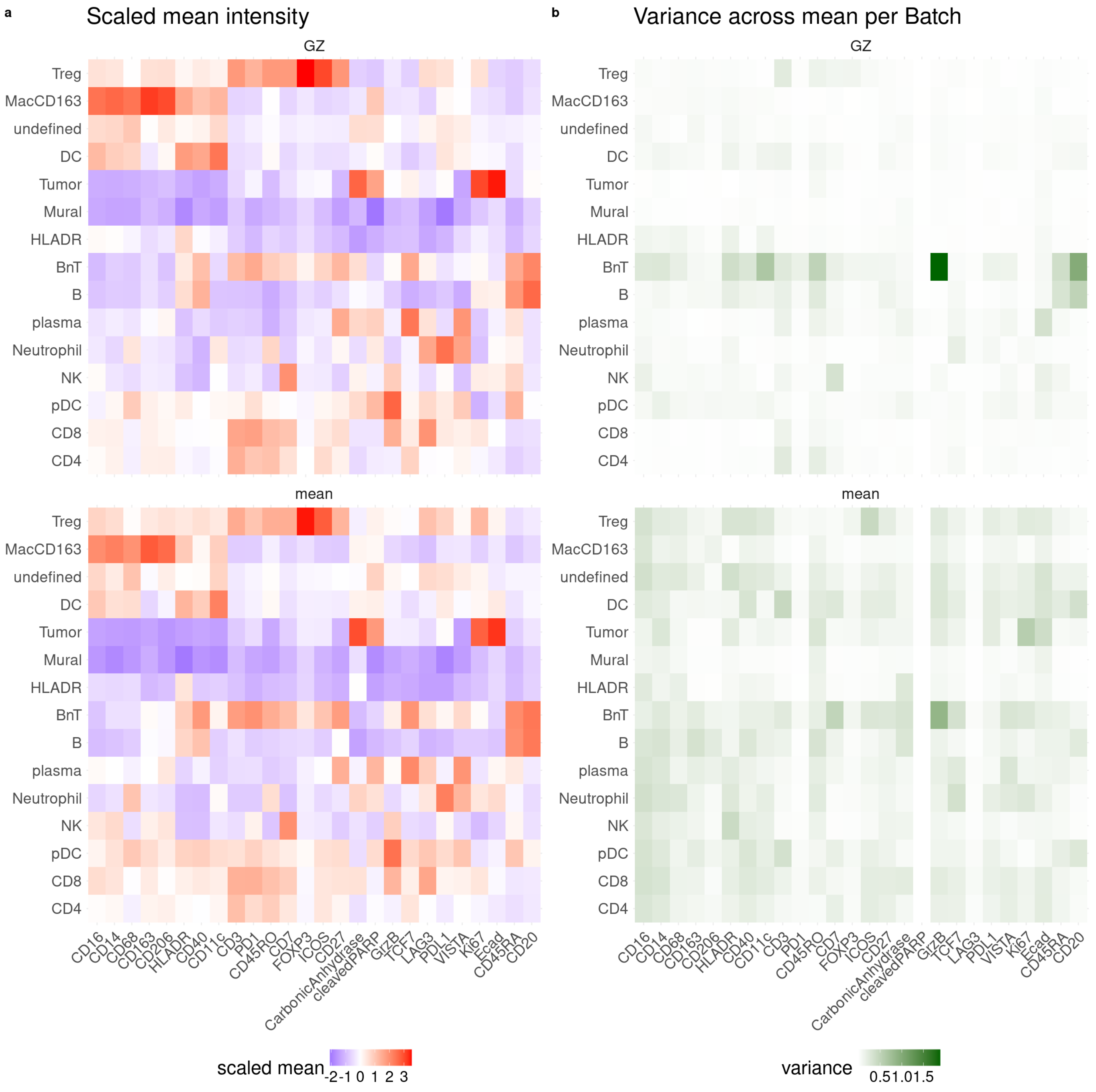


Supplementary Figure 6: Per cell type and marker mean intensities (a) and variances across batches (b) for the GZ normalized (top) and mean aggregated (bottom row) data.

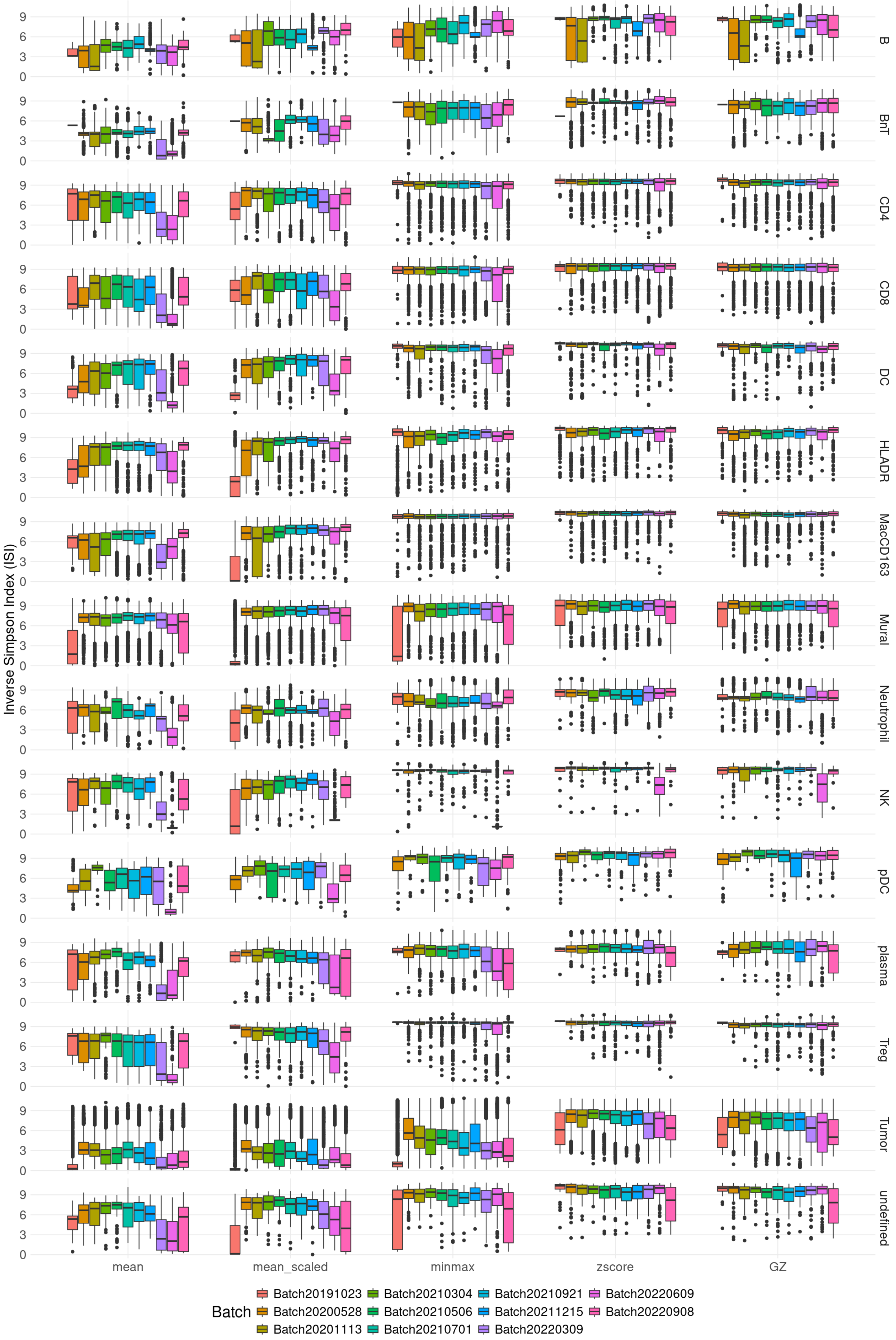


Supplementary Figure 7: Inverse Simpson Index (ISI) for the IMMUcan dataset split by cell type, batch and data normalization.

| normalization | ISI | NP | AMSP |
|---|---|---|---|
| mean | 3.381378 | 0.3245572 | 0.3865647 |
| GZ | 3.145048 | 0.3432475 | 0.2431093 |
| GZR | 3.251888 | 0.3419857 | 0.2521754 |
| minmax | 3.198622 | 0.3551953 | 0.1930311 |
| zscore | 3.150749 | 0.3417386 | 0.2401217 |
| mean_harmony | 3.106068 | 0.3614151 | 0.1775732 |
| mean_scaled_harmony | 3.034811 | 0.3752724 | 0.1573694 |
| GZ_harmony | 3.456902 | 0.2880463 | 0.2768690 |
| minmax_harmony | 3.320926 | 0.3066338 | 0.2522143 |
| zscore_harmony | 3.397251 | 0.2905644 | 0.2821073 |

Supplementary Table 1: Celltype annotation metrics in the IMMUcan dataset for different normalizations. ISI: inverse simpson index, NP: Neighborhood Purity, AMSP: the Adjusted Mean Shortest Path.

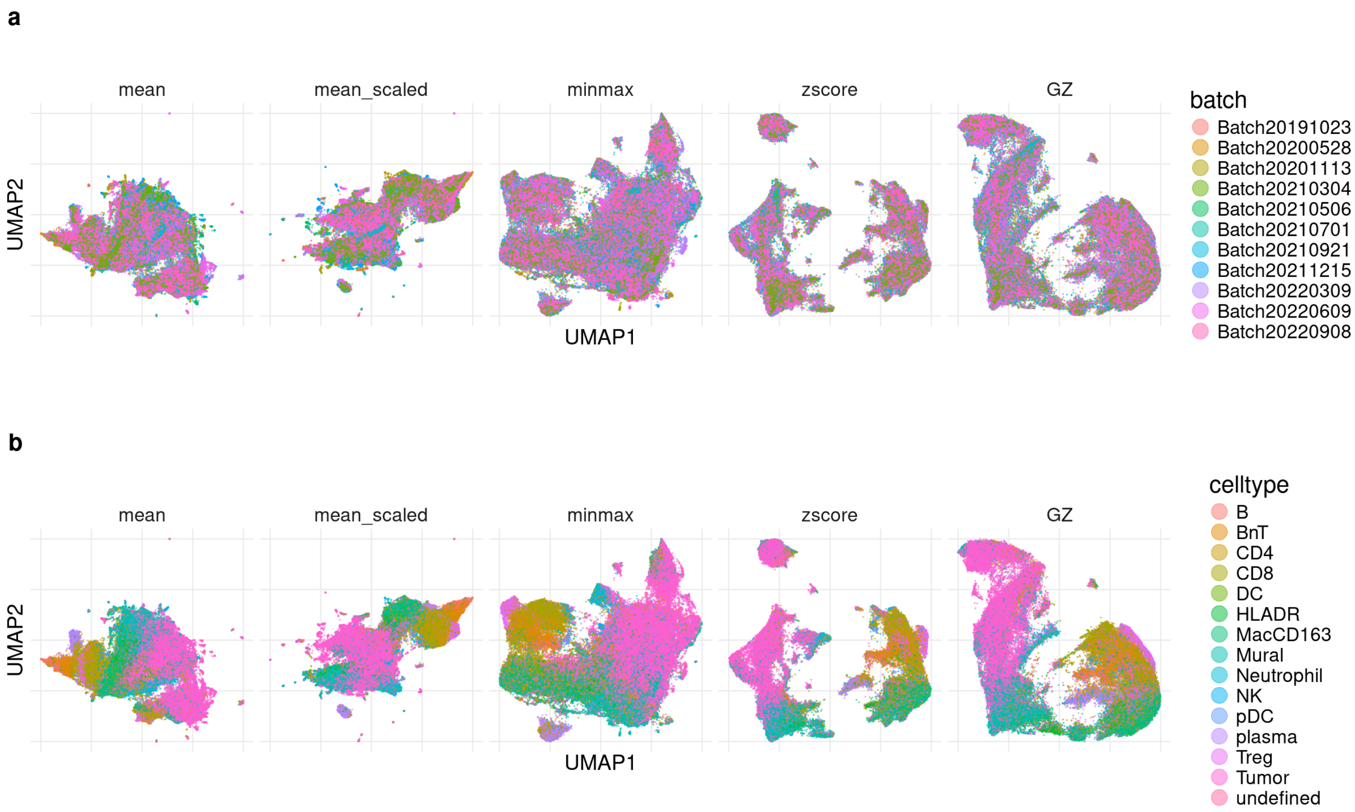


Supplementary Figure 8: UMAP for five different normalizations followed by harmony integration at the cell level in IMMUcan dataset: 'mean' (no normalization), 'mean_scaled' (minmax scaling at cell level), 'minmax' (per ROI min-max normalization), 'zscore' (per ROI z-score normalization), 'GZ' (per ROI GZ normalization). a) colored by batch. b) colored by celltype.

manuscript_files/figure-pdf/figsup-44-1.png

Supplementary Figure 9: AUROC for 'GZ' versus 'mean' for different markers and cell types.

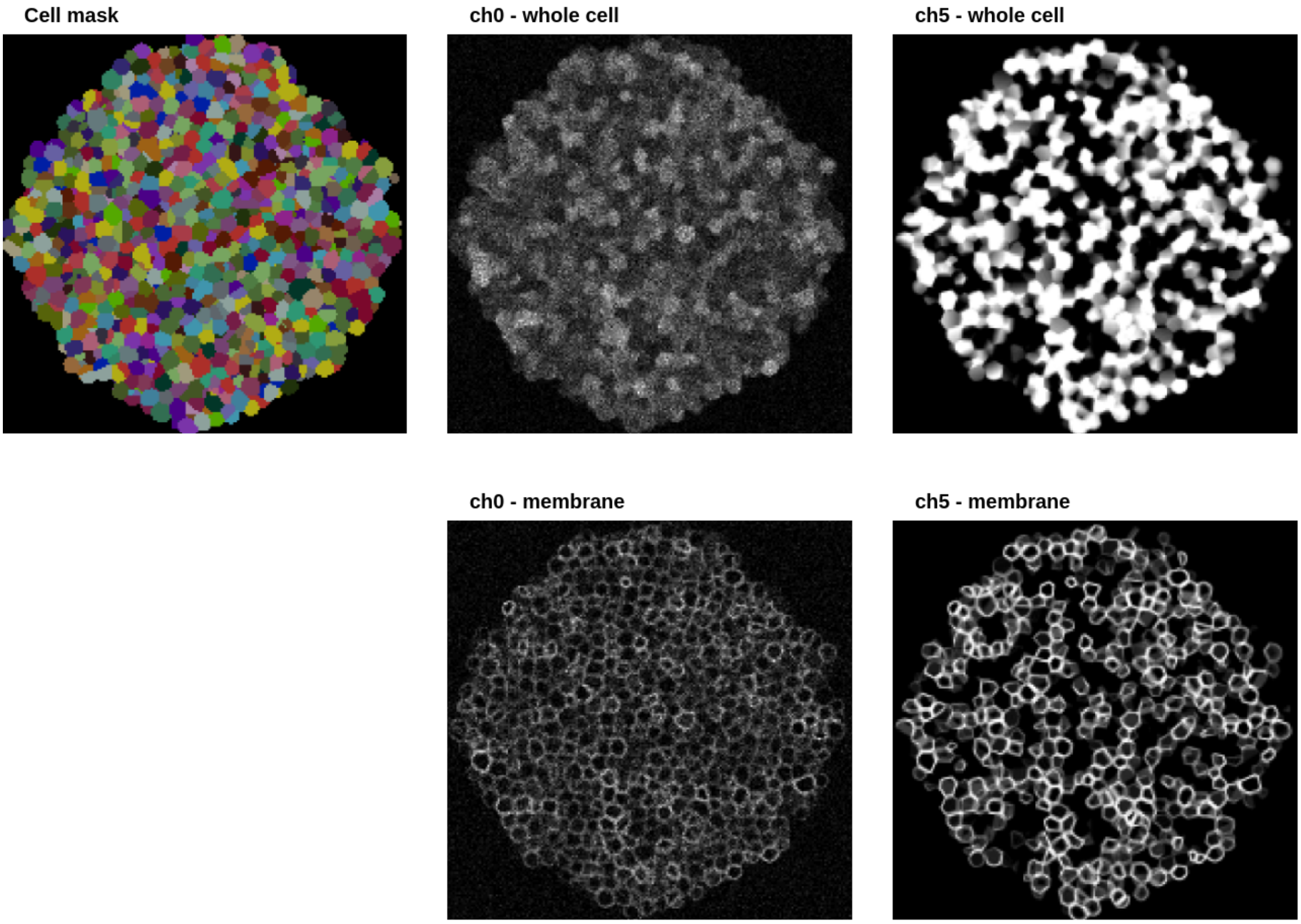


Supplementary Figure 10: Simulated data example image. Left: Cell Mask. Middle top: intensity whole cell for noisy channel ch0. Middle bottom: intensity membrane for noisy channel ch0. Right top: intensity whole cell for no noise channel ch5. Right bottom: intensity membrane for no noise channel ch5.

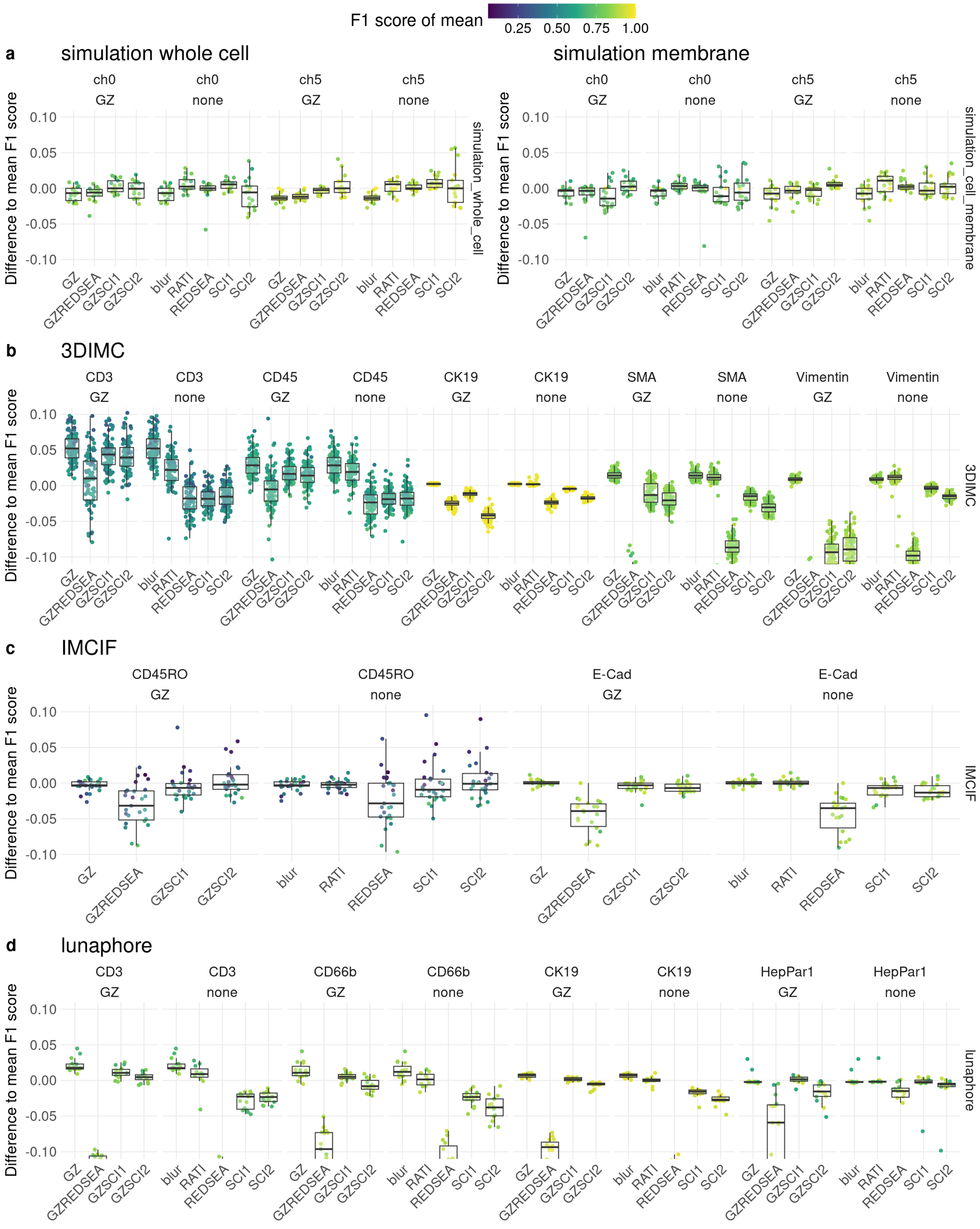


Supplementary Figure 11: Relative F1 scores (difference to baseline) for data set for different methods, normalization and markers.

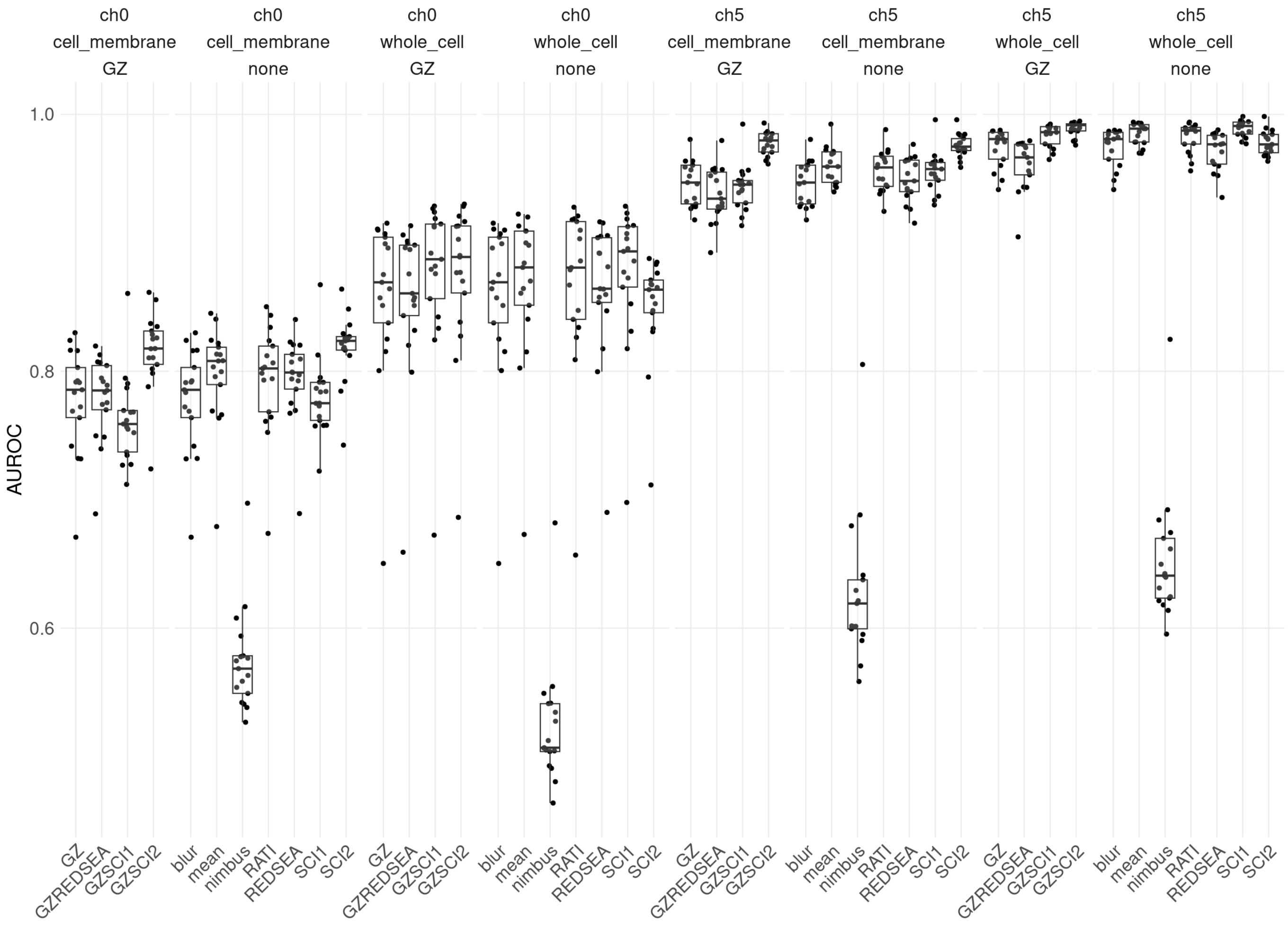


Supplementary Figure 12: AUROC for simulated data set for different methods, normalization and markers.

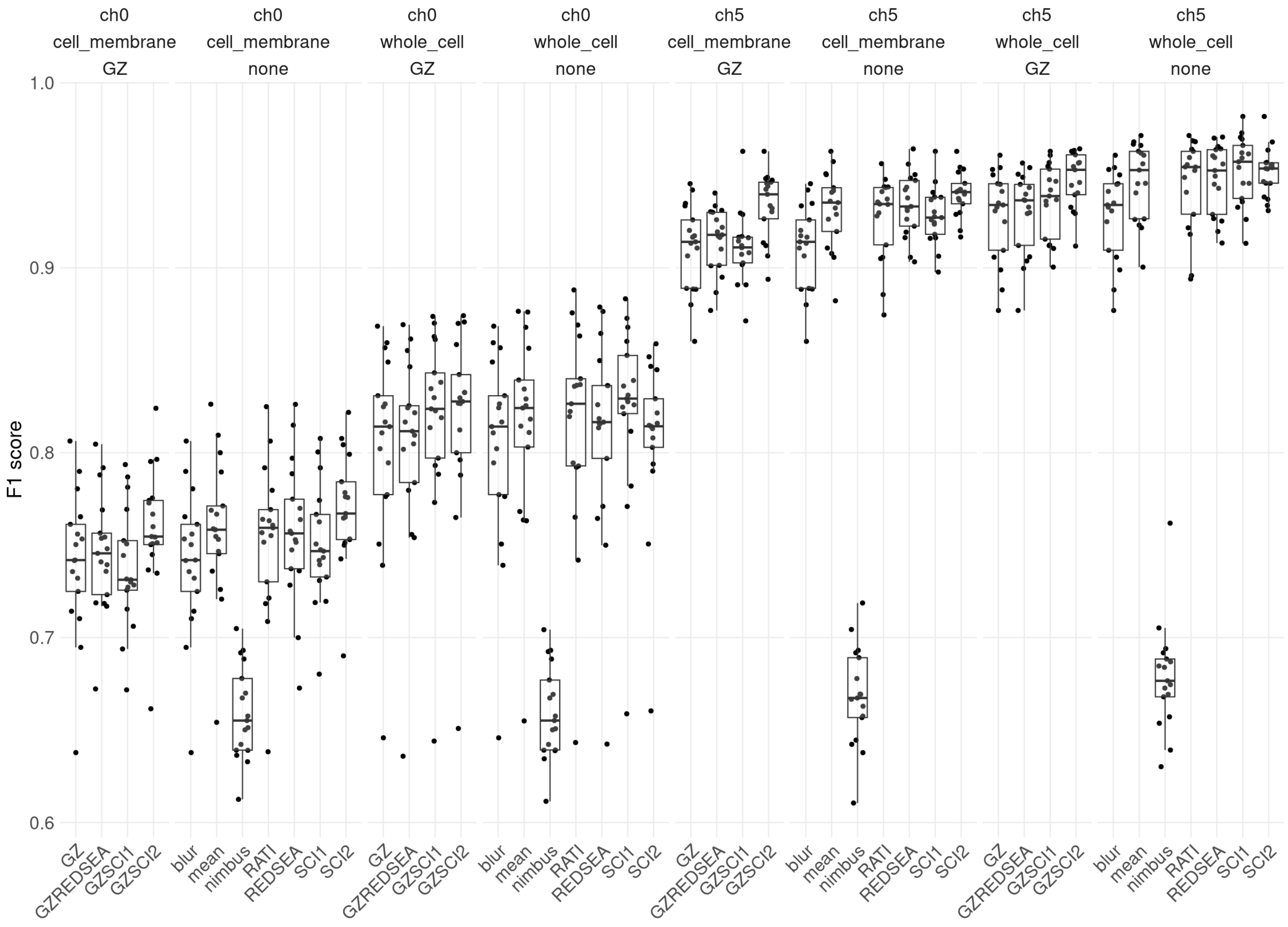


Supplementary Figure 13: F1 scores for simulated data set for different methods, normalization and markers.

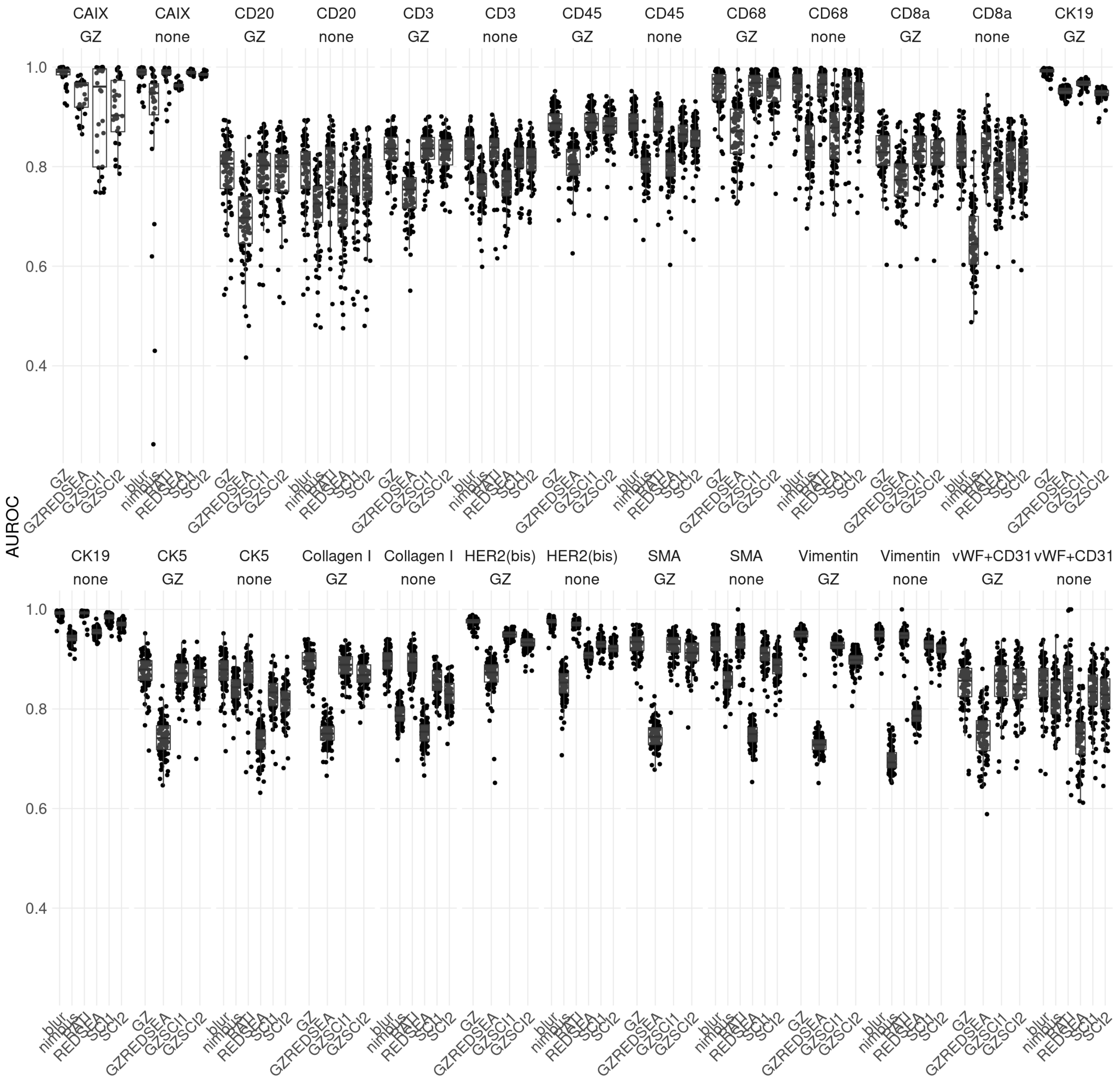


Supplementary Figure 14: AUROC for ‘Second’ 3DIMC for different methods, normalization and markers.

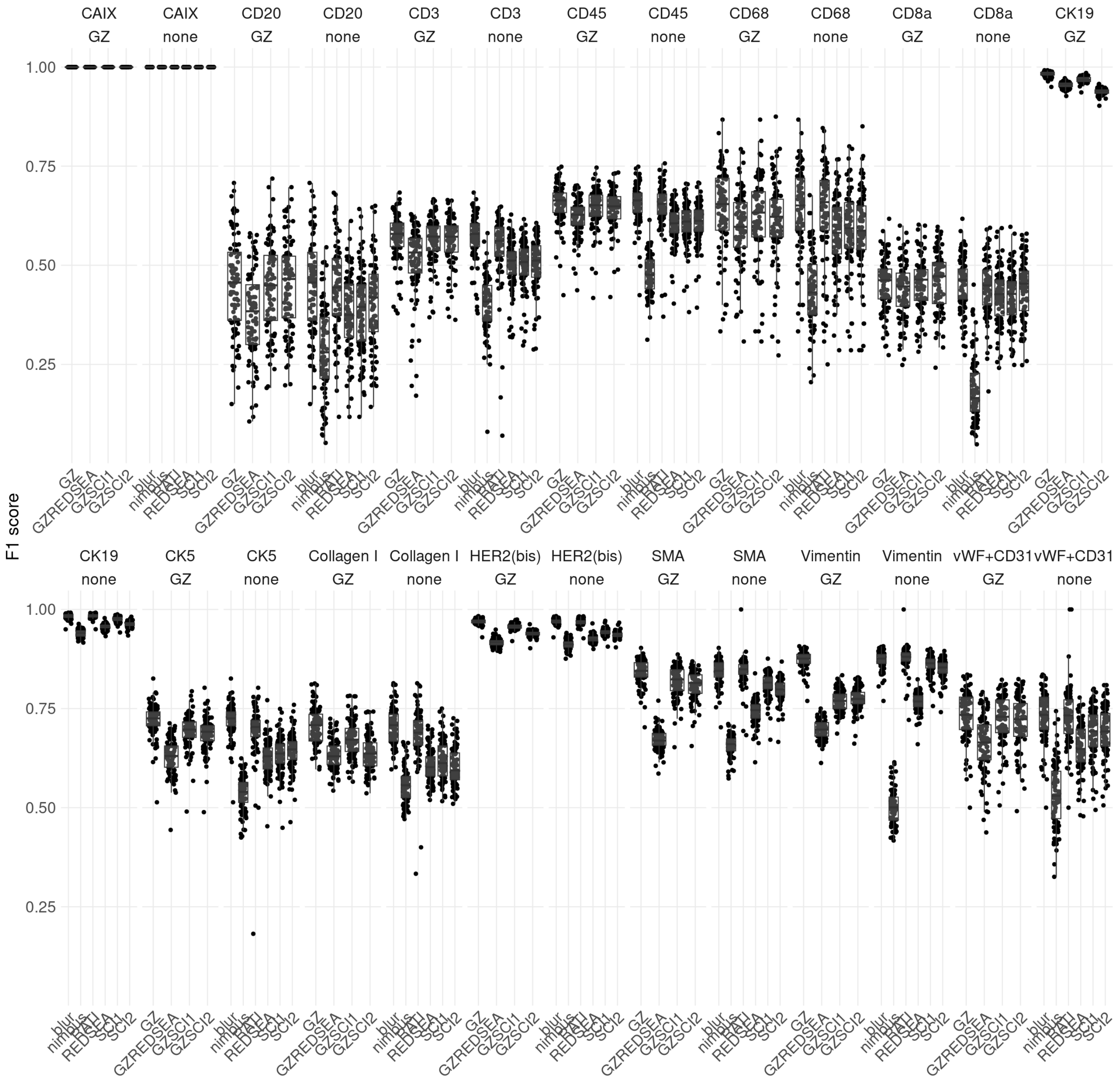


Supplementary Figure 15: F1 scores for ‘Second’ 3DIMC for different methods, normalization and markers.

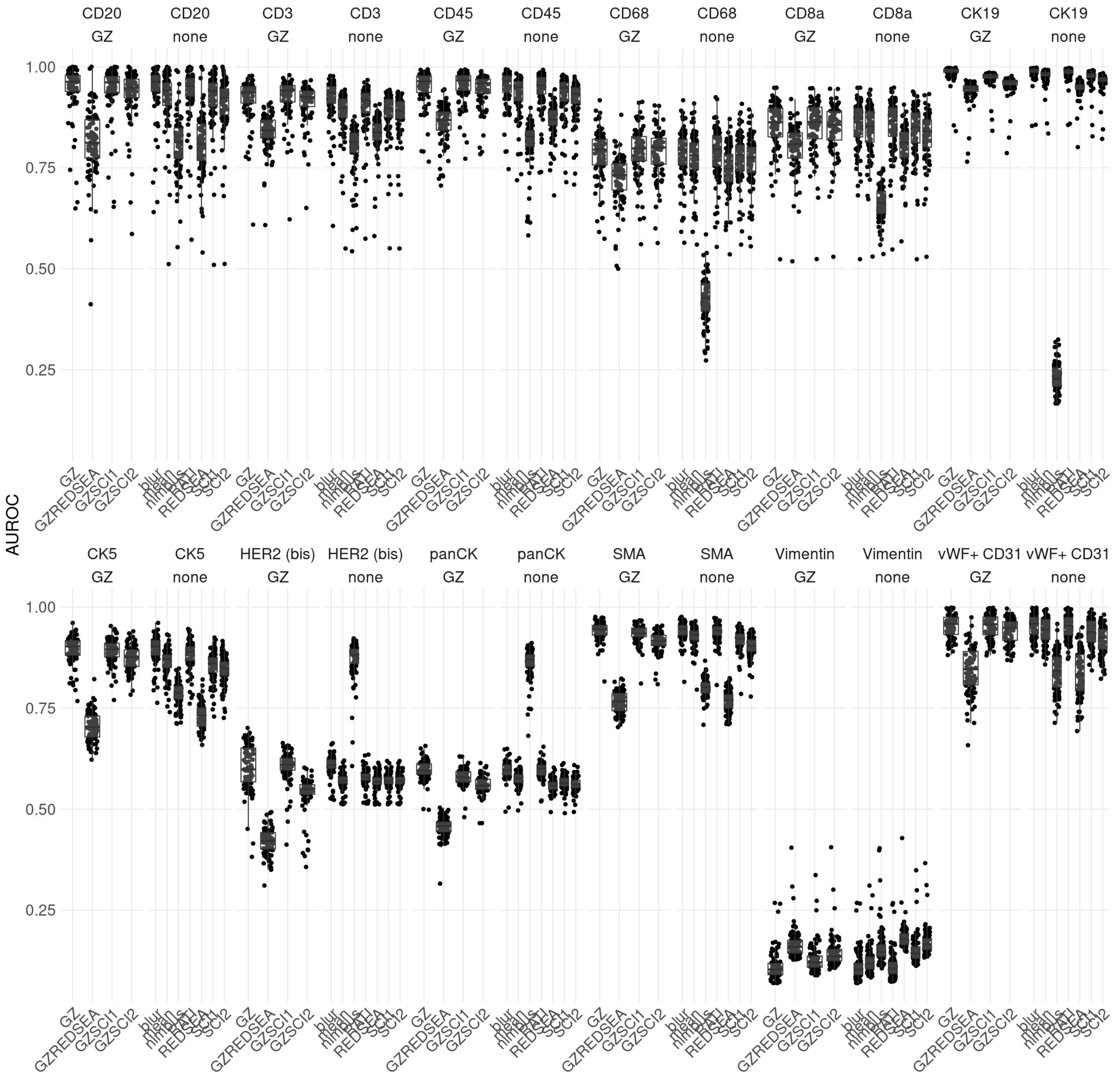


Supplementary Figure 16: AUROC for 'Main' 3DIMC for different methods, normalization and markers.

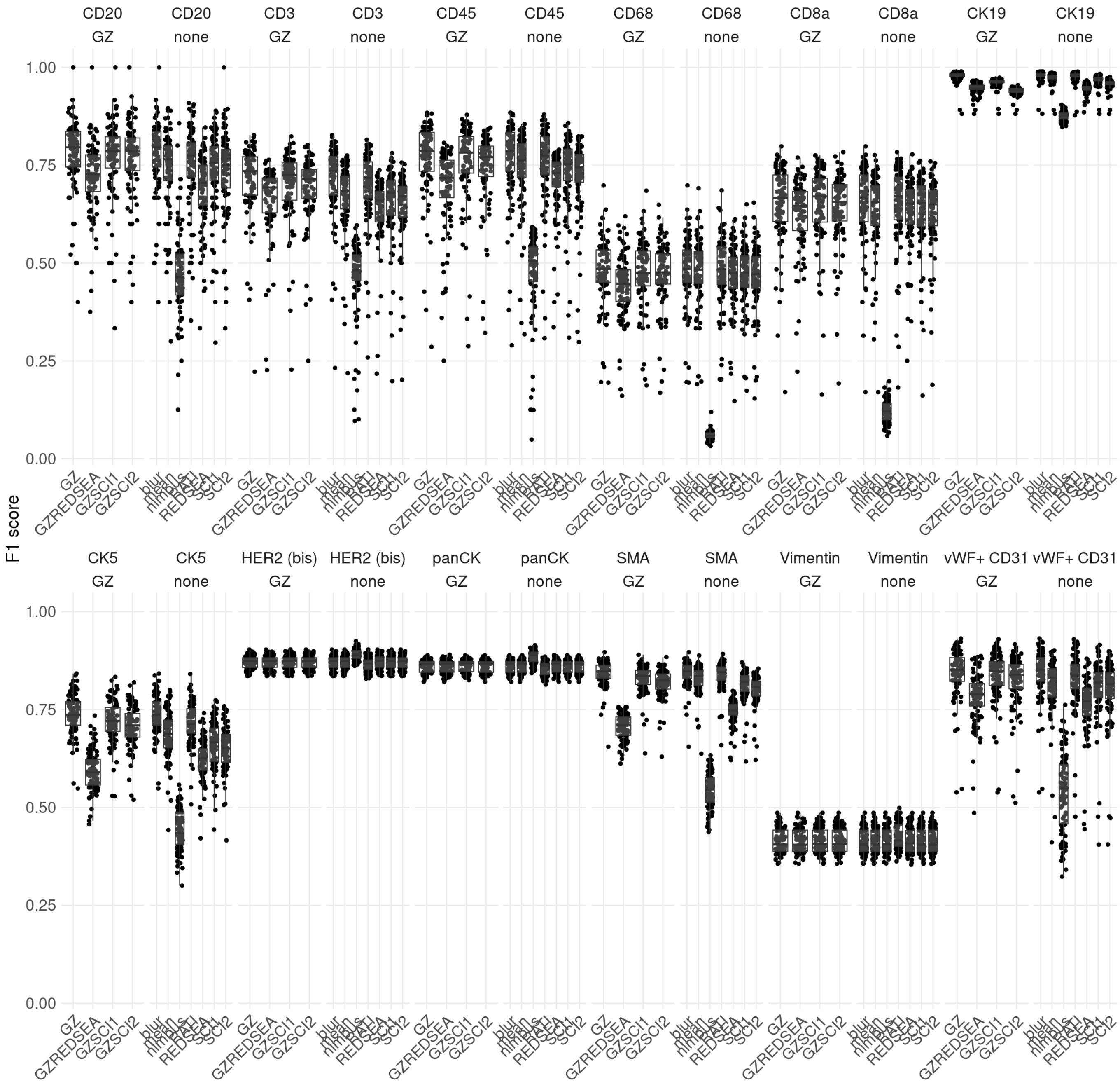


Supplementary Figure 17: F1 scores for 'Main' 3DIMC for different methods, normalization and markers.

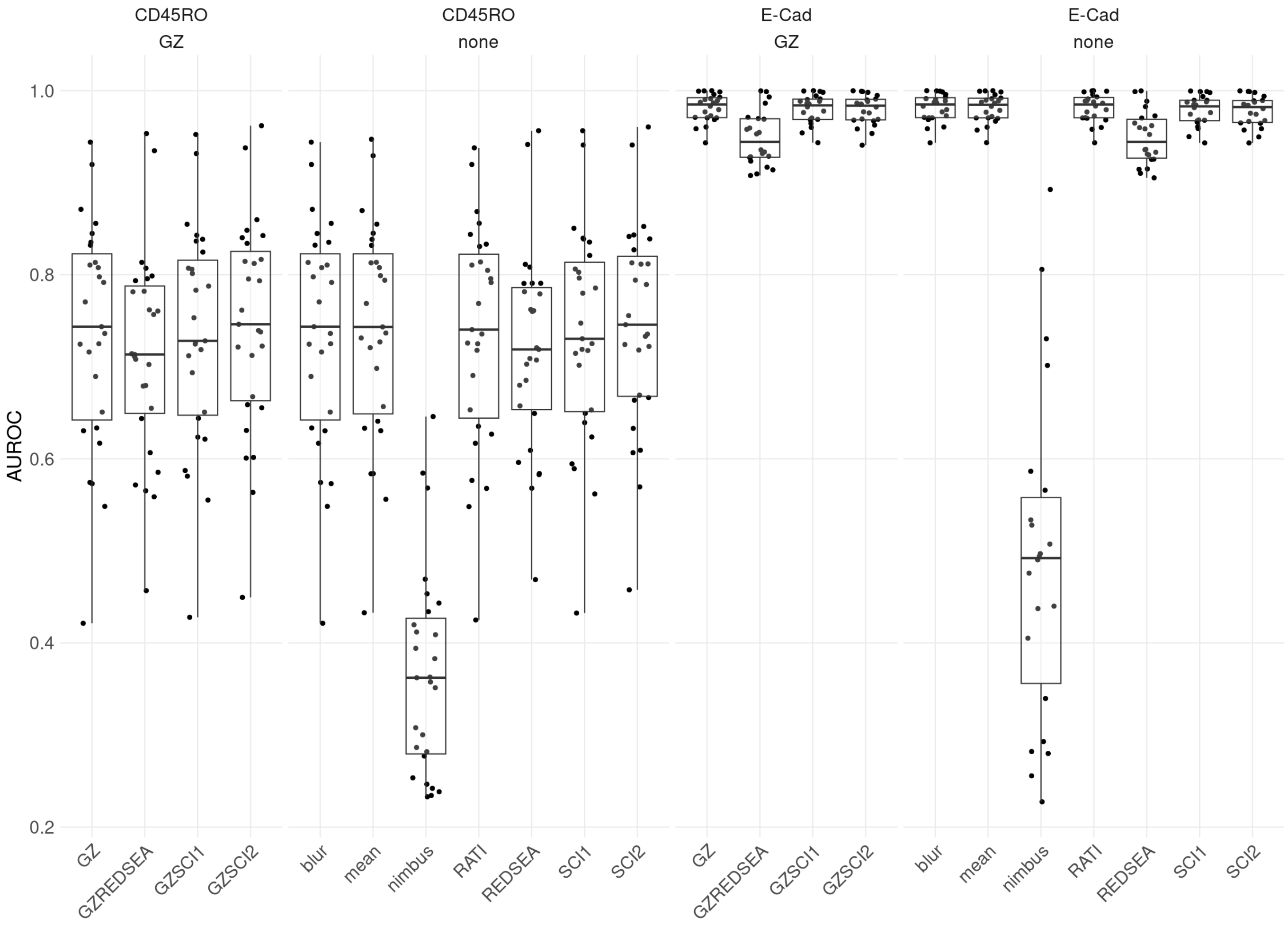


Supplementary Figure 18: AUROC for IMCIF data set for different methods, normalization and markers.

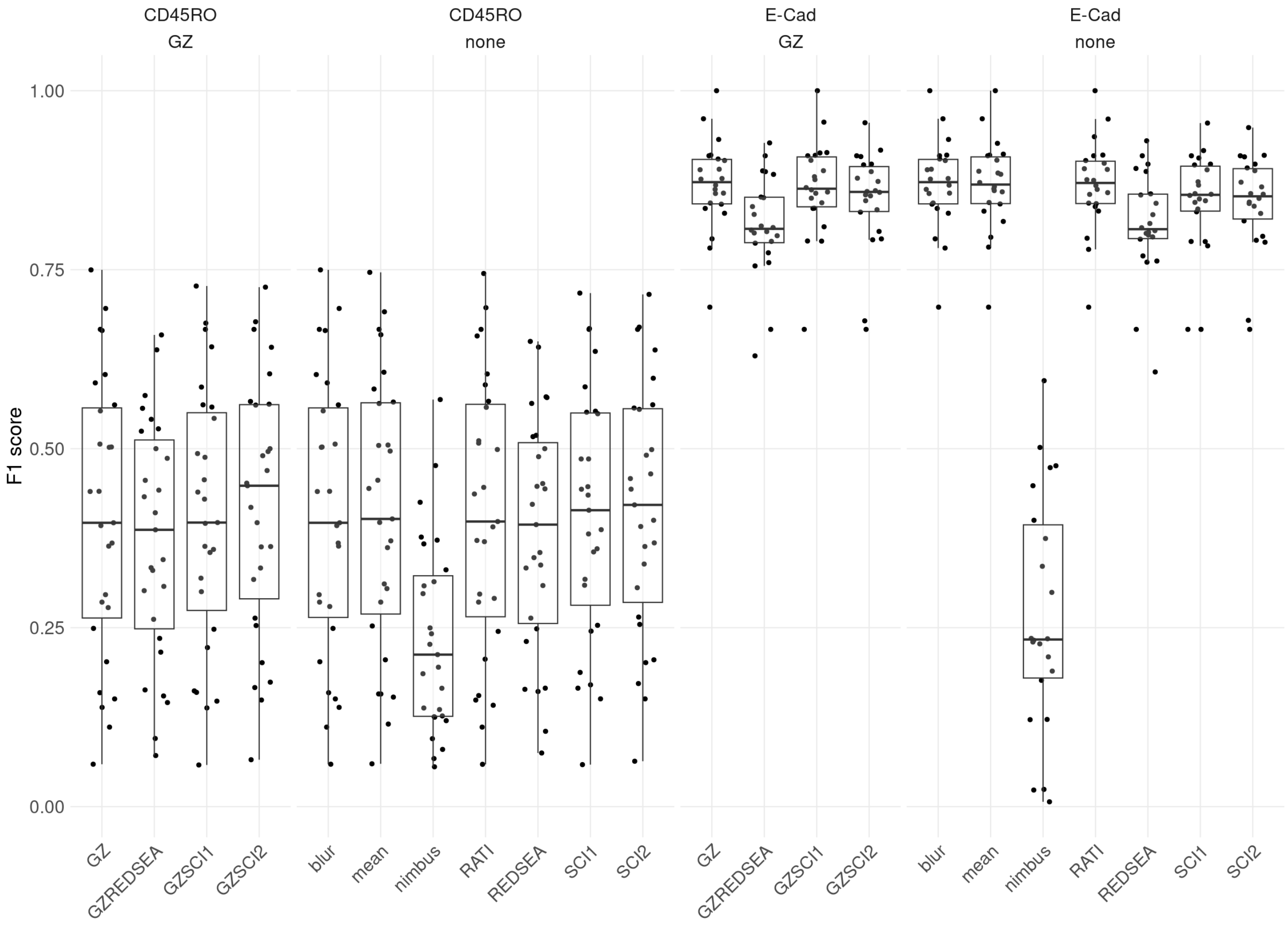


Supplementary Figure 19: F1 scores for IMCIF data set for different methods, normalization and markers.

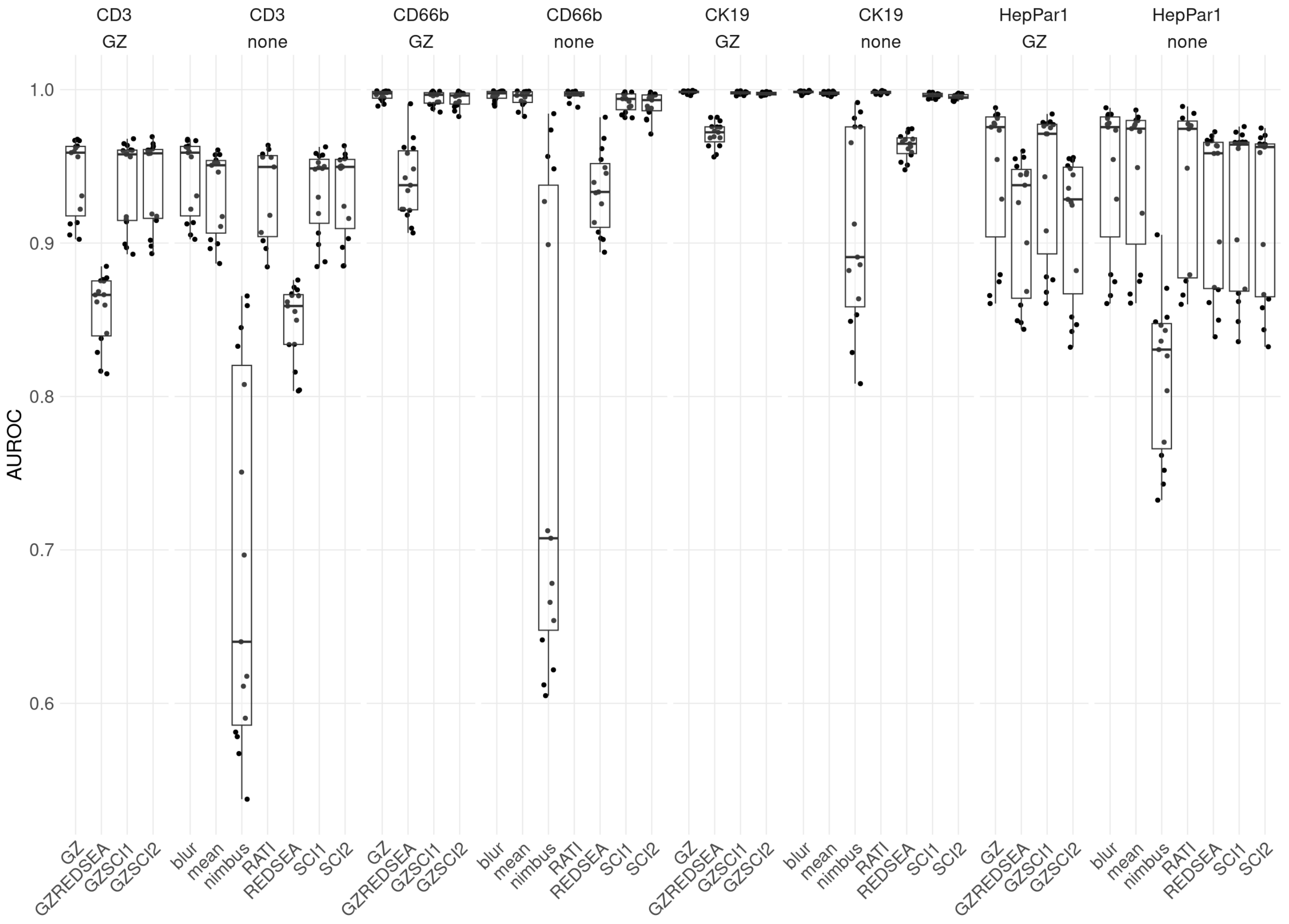


Supplementary Figure 20: AUROC for lunaphore data set for different methods, normalization and markers.

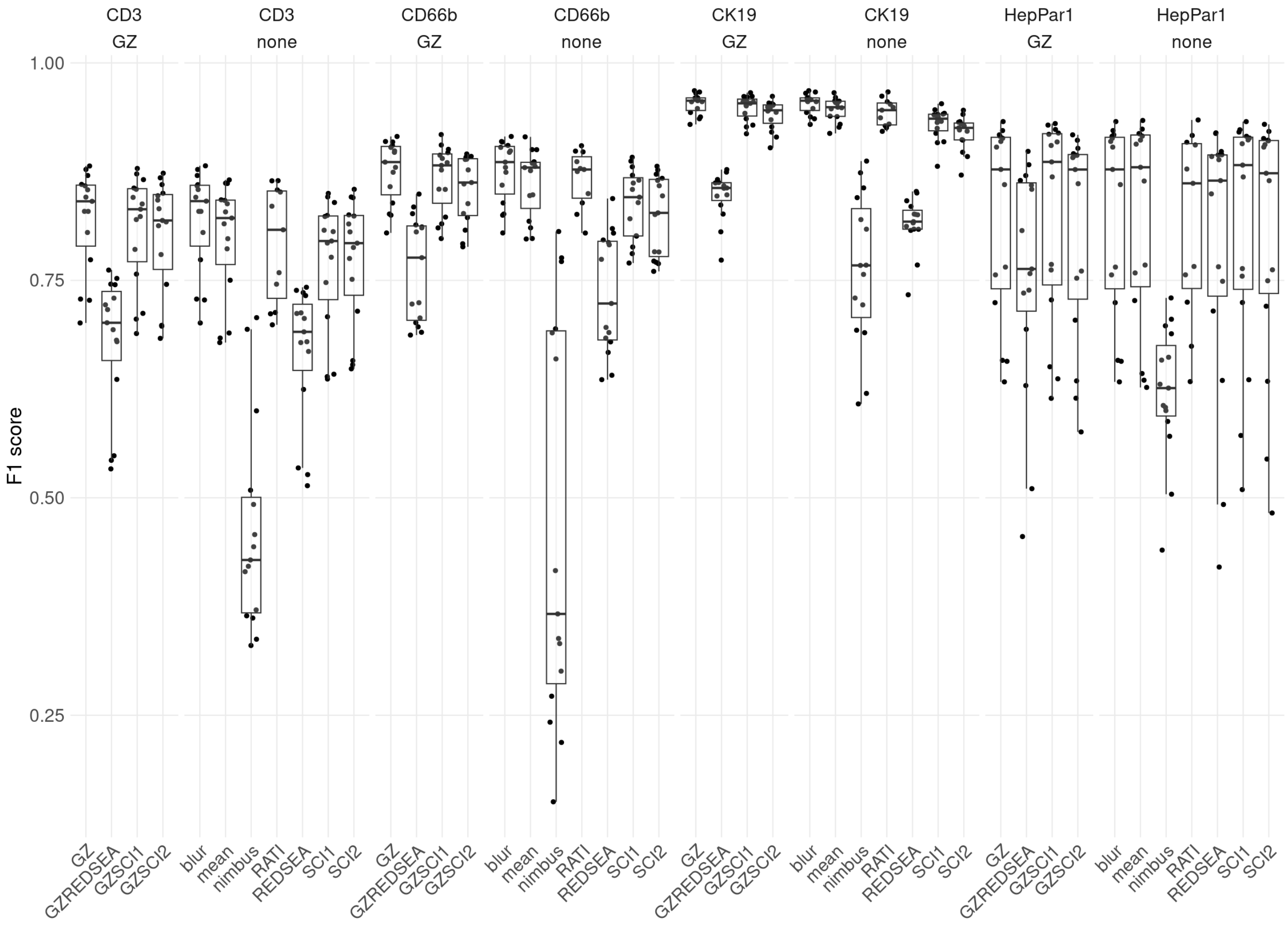


Supplementary Figure 21: F1 score for lunaphore data set for different methods, normalization and markers.

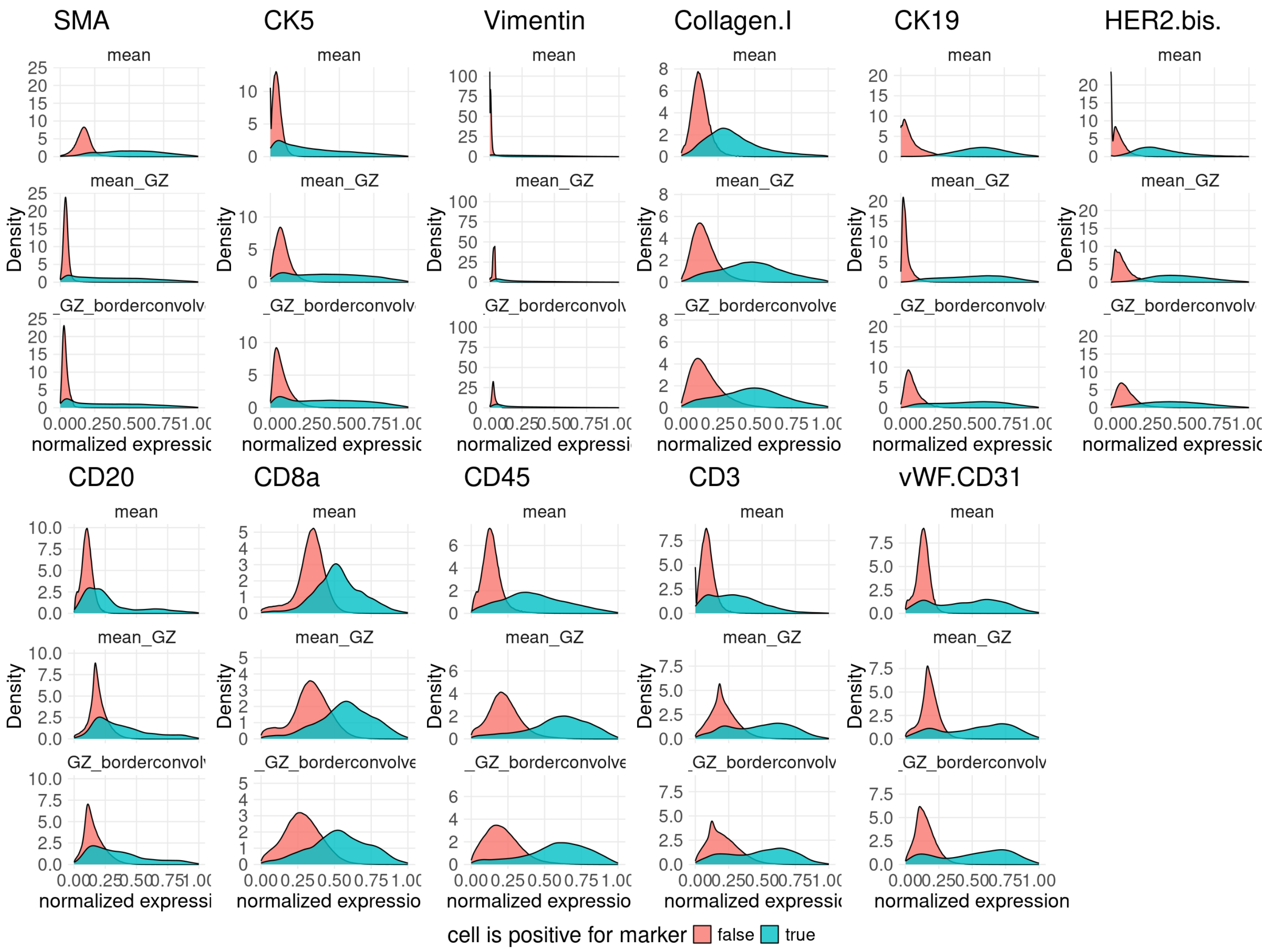


Supplementary Figure 22: Per marker and normalization distribution of cell intensities for 3D IMC dataset colored by marker positivity."

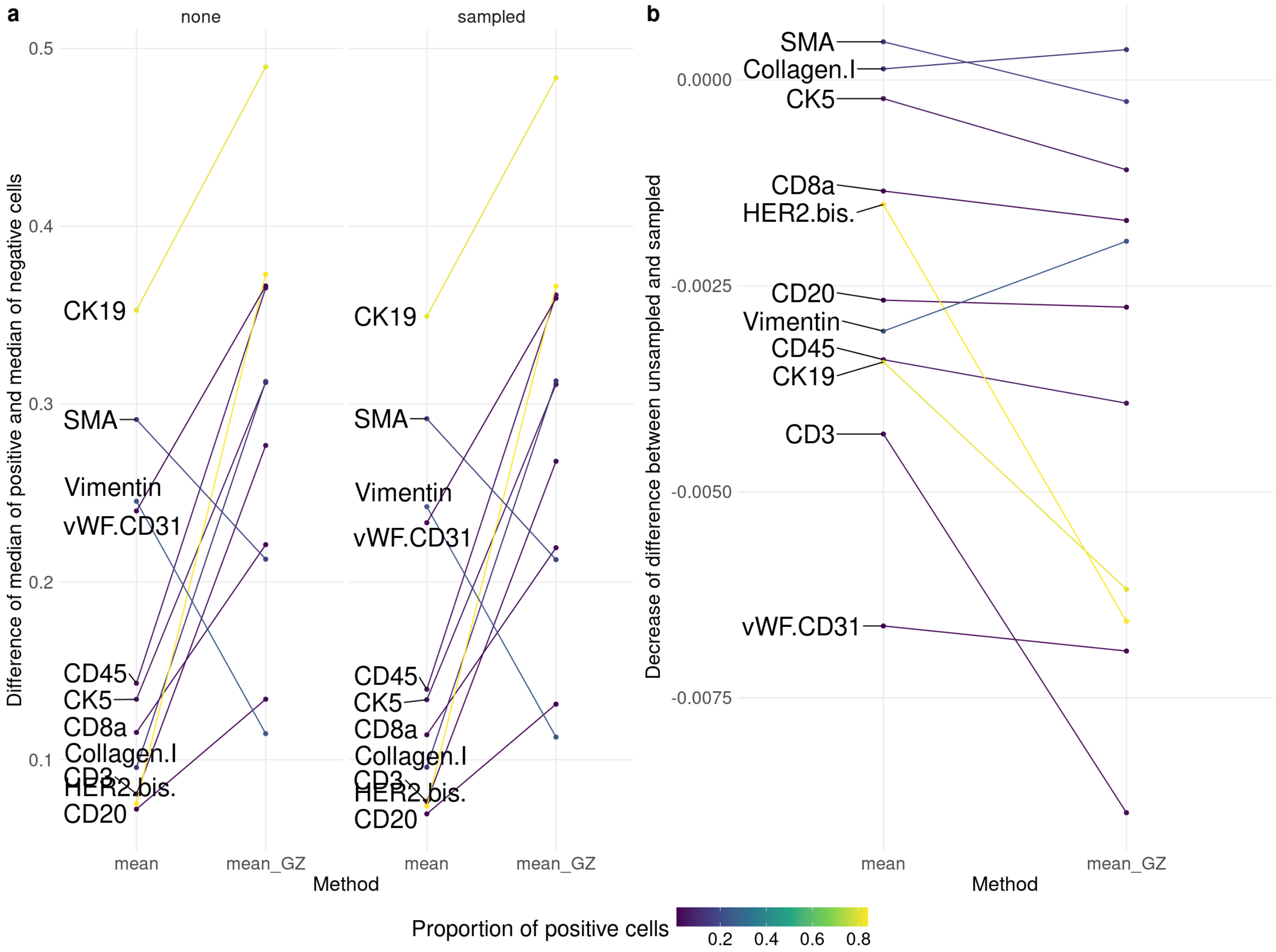


a) Difference between marker positive and marker negative populations for different normalizations for correct segmentation mask (left) and spillover simulated segmentation mask (right). b) Difference between a) left and a) right.

Supplementary Figure 23

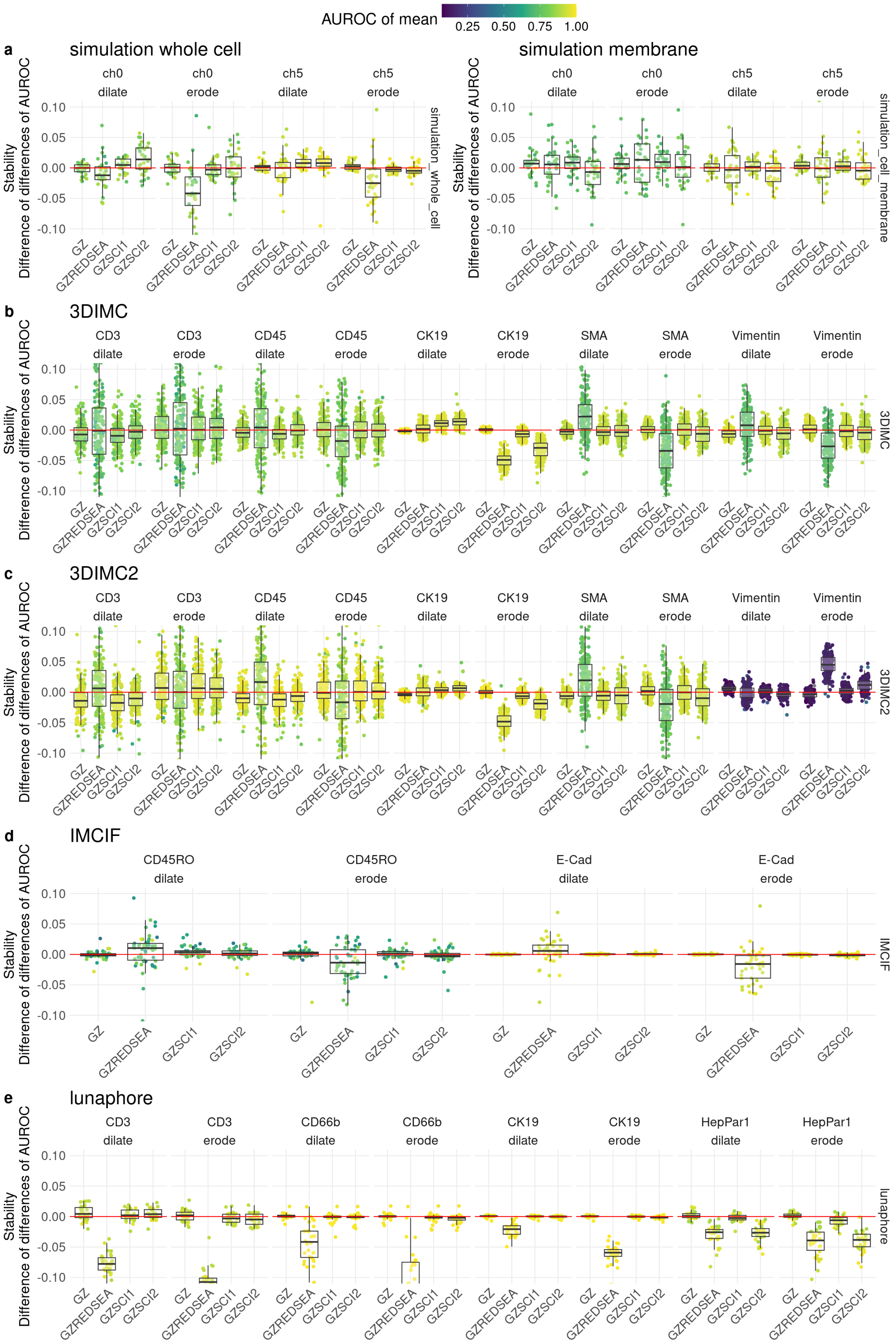


Supplementary Figure 24: Difference score calculated by first normalizing the AUROC by dividing by the baseline (mean aggregation) for both correct segmentation and spillover simulated segmentation, followed by taking the difference between the two. Values larger then zero means spillover simulated values are higher than for the baseline and thus more stable.

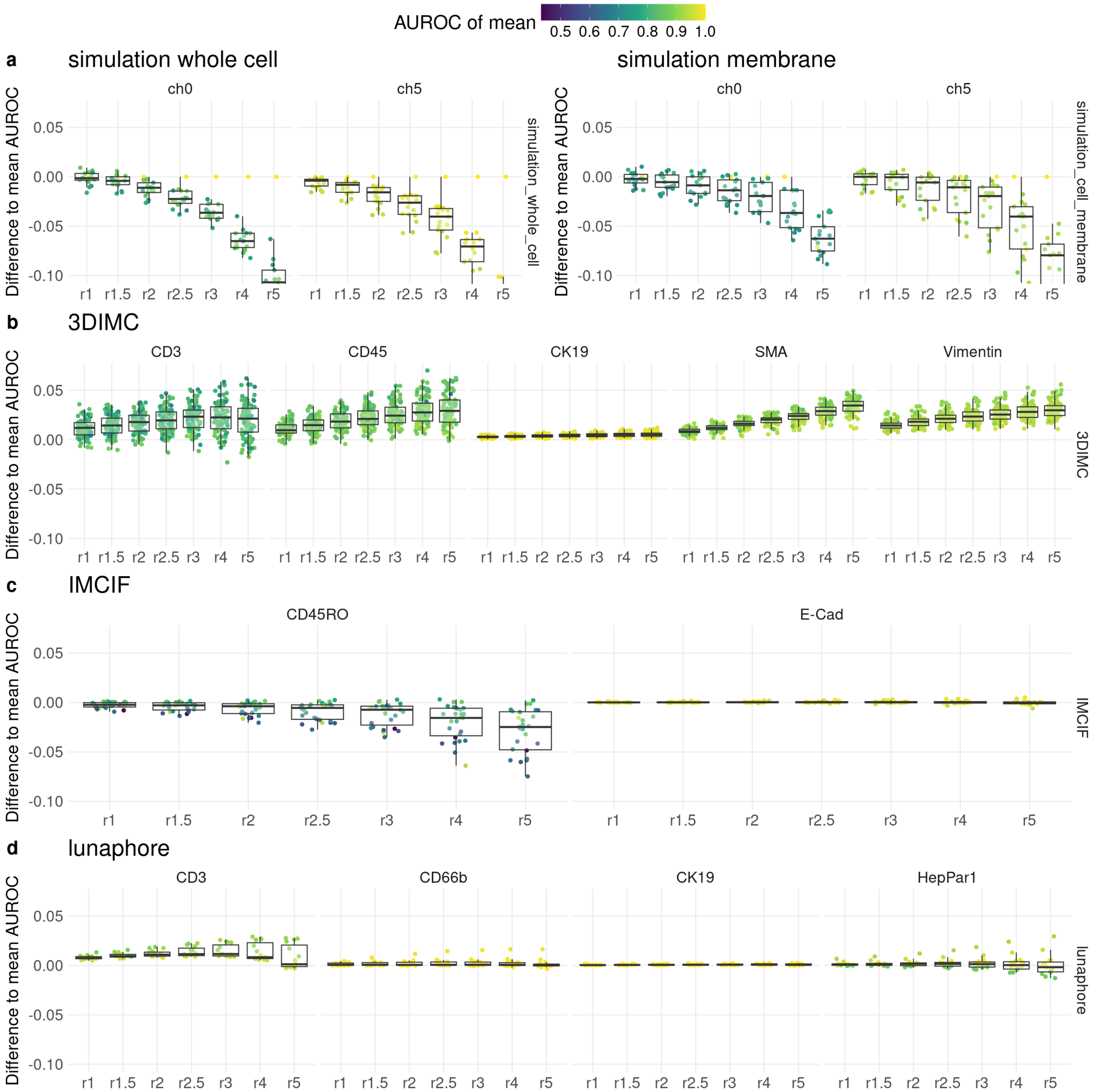


Supplementary Figure 25: Effect of neighborhood size in terms of distance from center pixel.

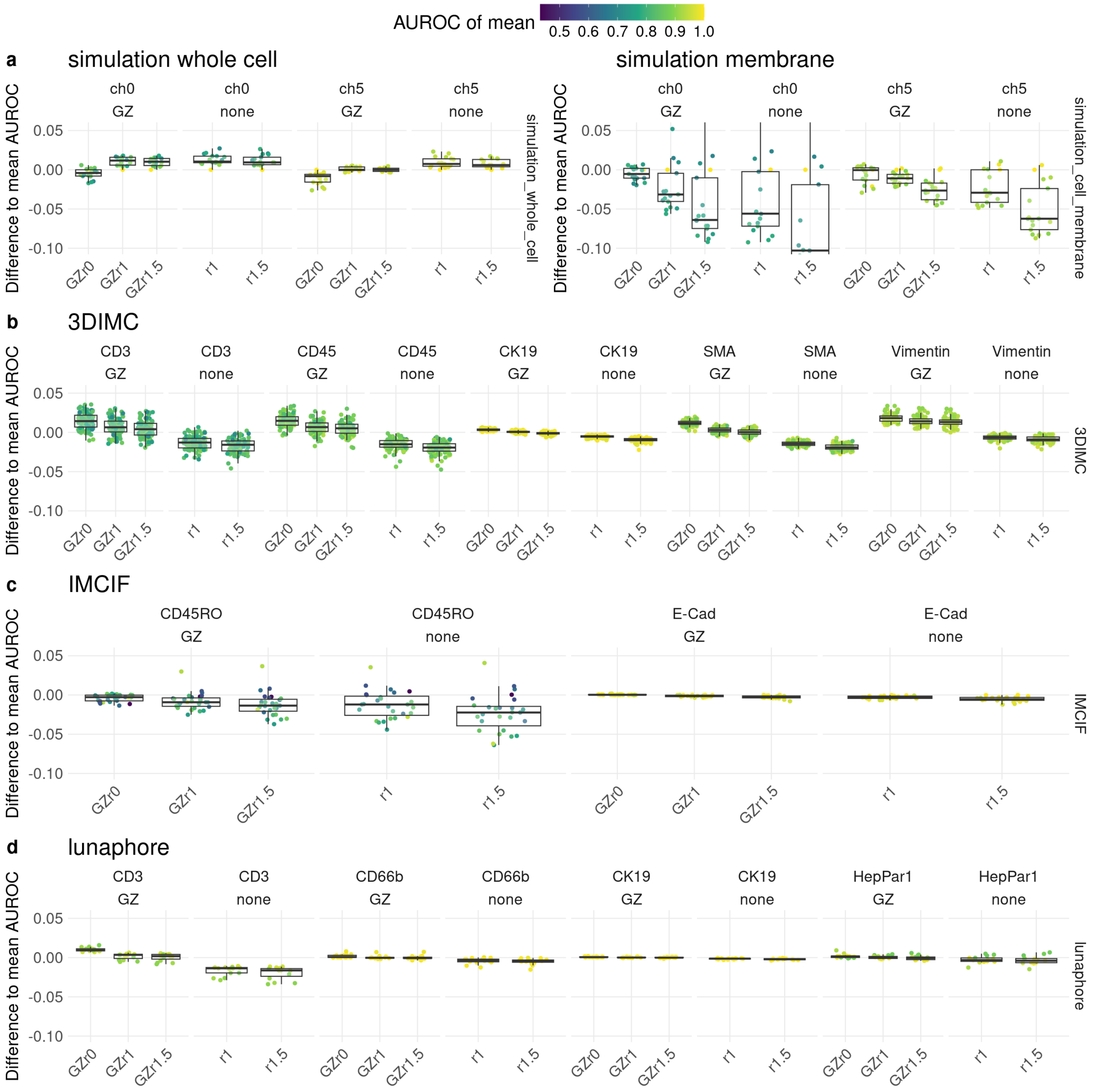


Supplementary Figure 26: Effect of cell mask erosion, r is the radius of erosion: r0 no erosion, r1 $3 \times 3$ cross kernel, r1.5 $3 \times 3$ rectangular kernel.

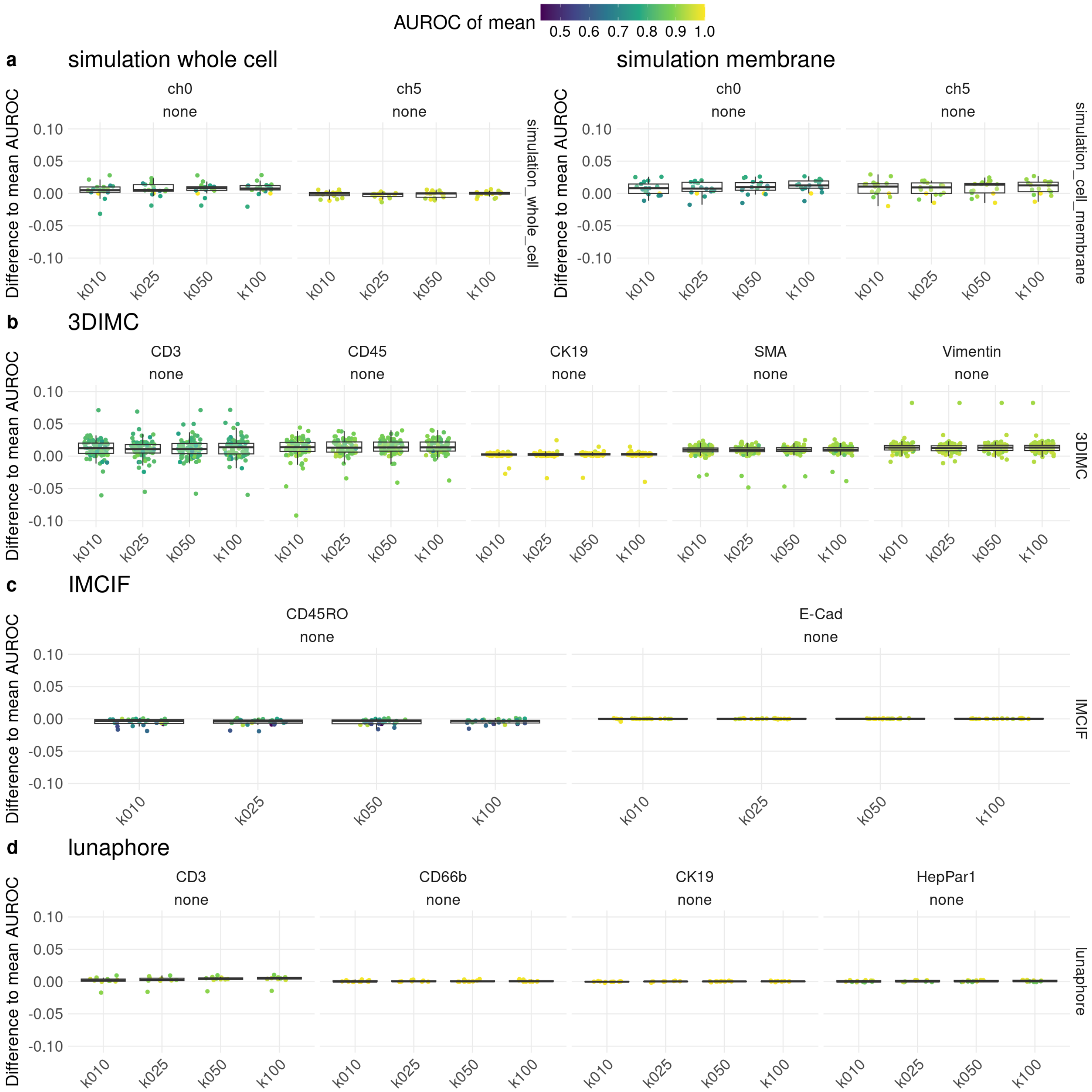


Supplementary Figure 27: Effect of number of iterations in bootstrapping k.

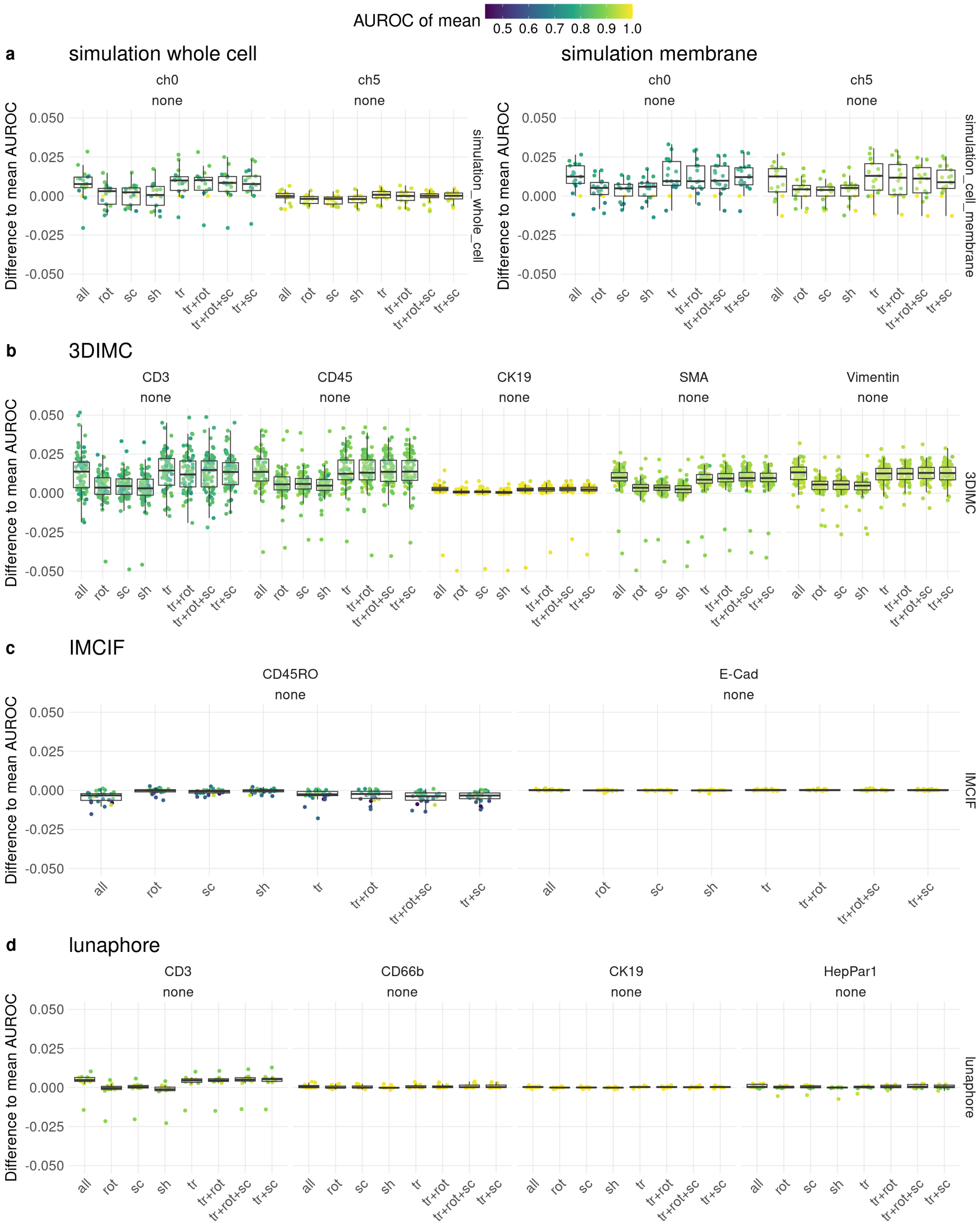


Supplementary Figure 28: Effect of which parameters of the affine transform are sampled. Only specified parameters are sampled, the other are kept at identity. tr: translation, rot: rotation, sc: scaling, sh: shearing.

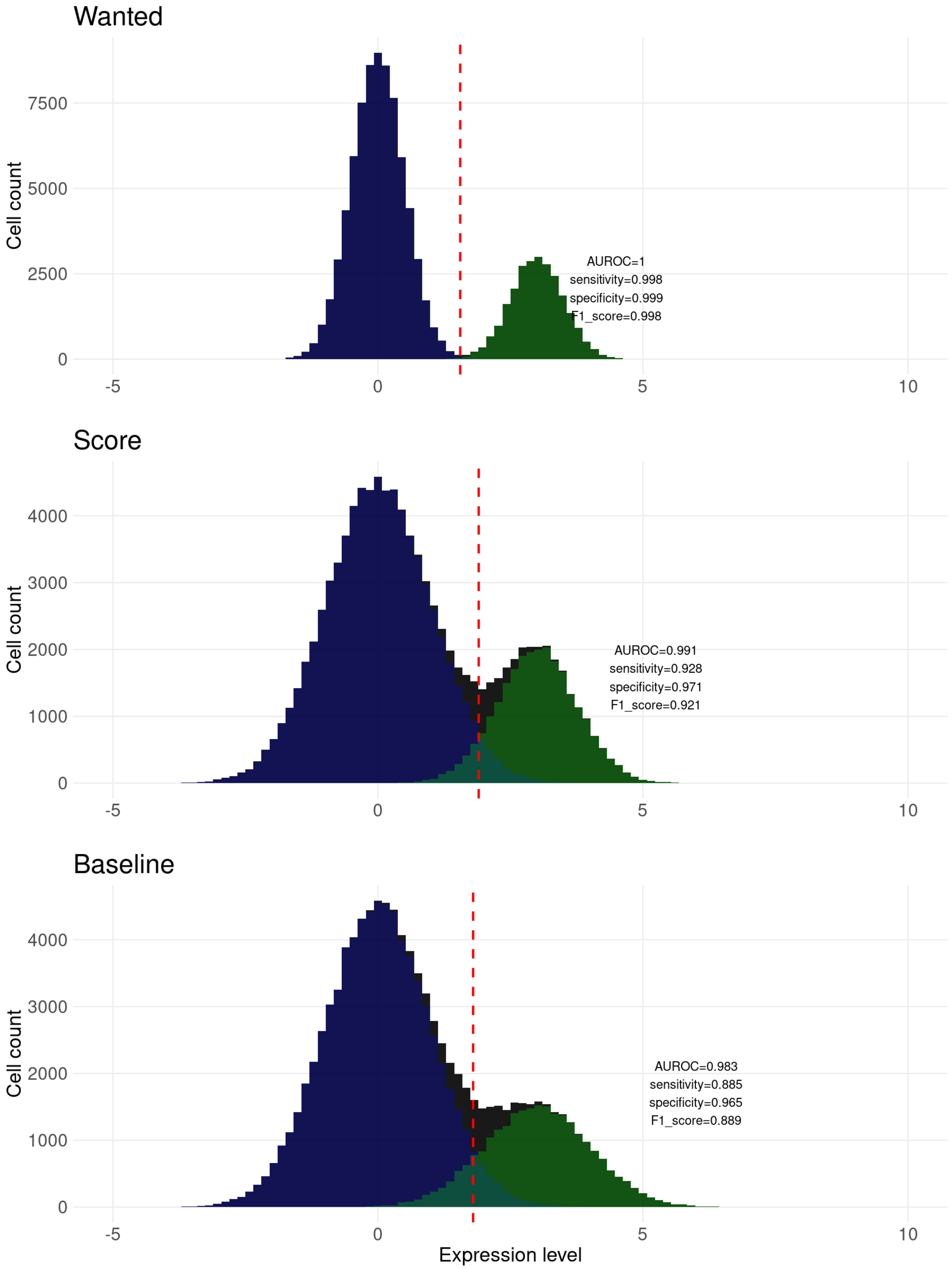


Supplementary Figure 29: Example simulated data showing behaviour of AUROC and F1-score on separation of positive and negative cells for three scenarios: Wanted, Score, and Baseline.

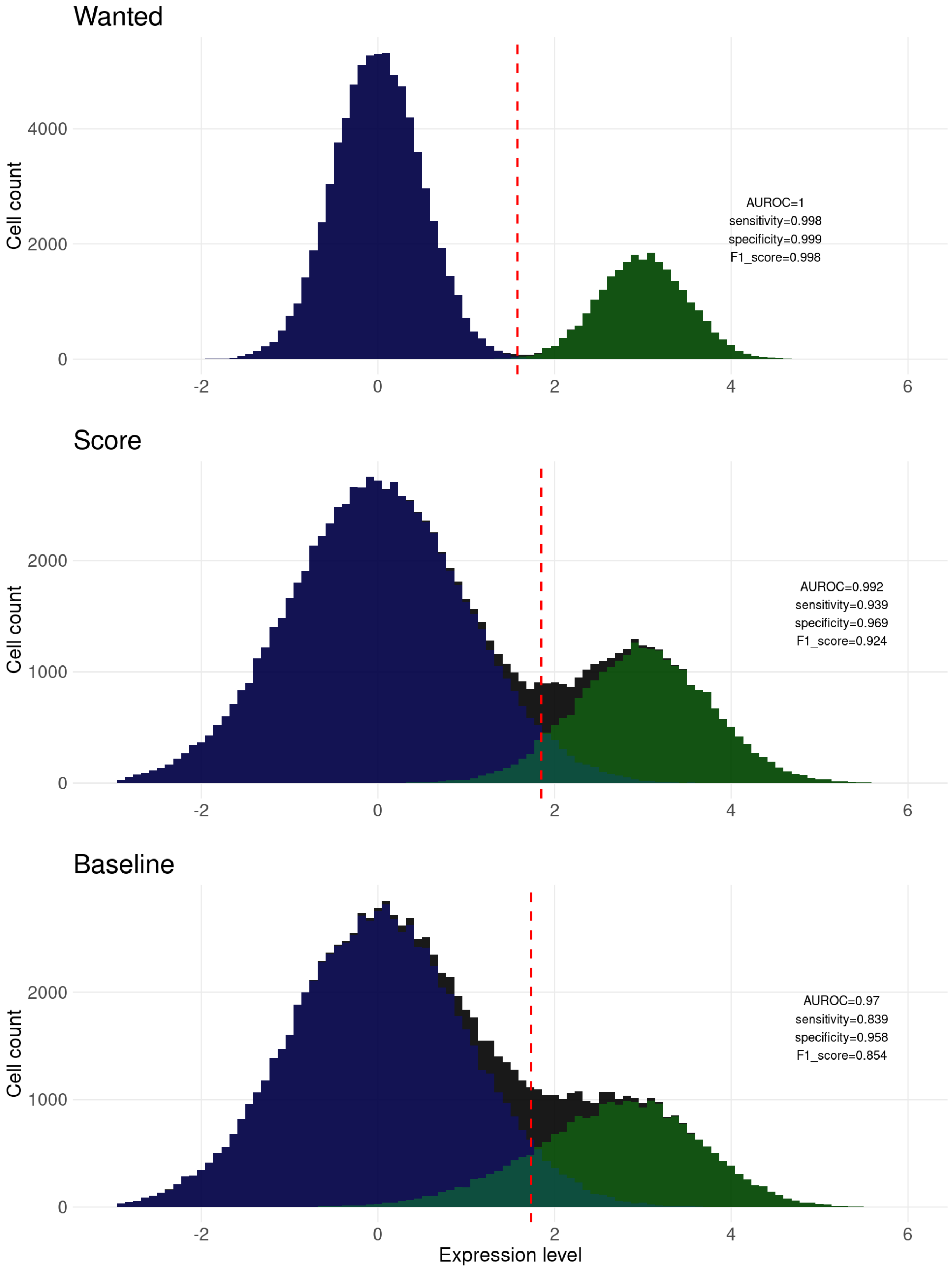


Supplementary Figure 30: Example simulated data showing behaviour of AUROC and F1-score on separation of positive and negative cells for three scenarios: Wanted, Score, and Baseline.

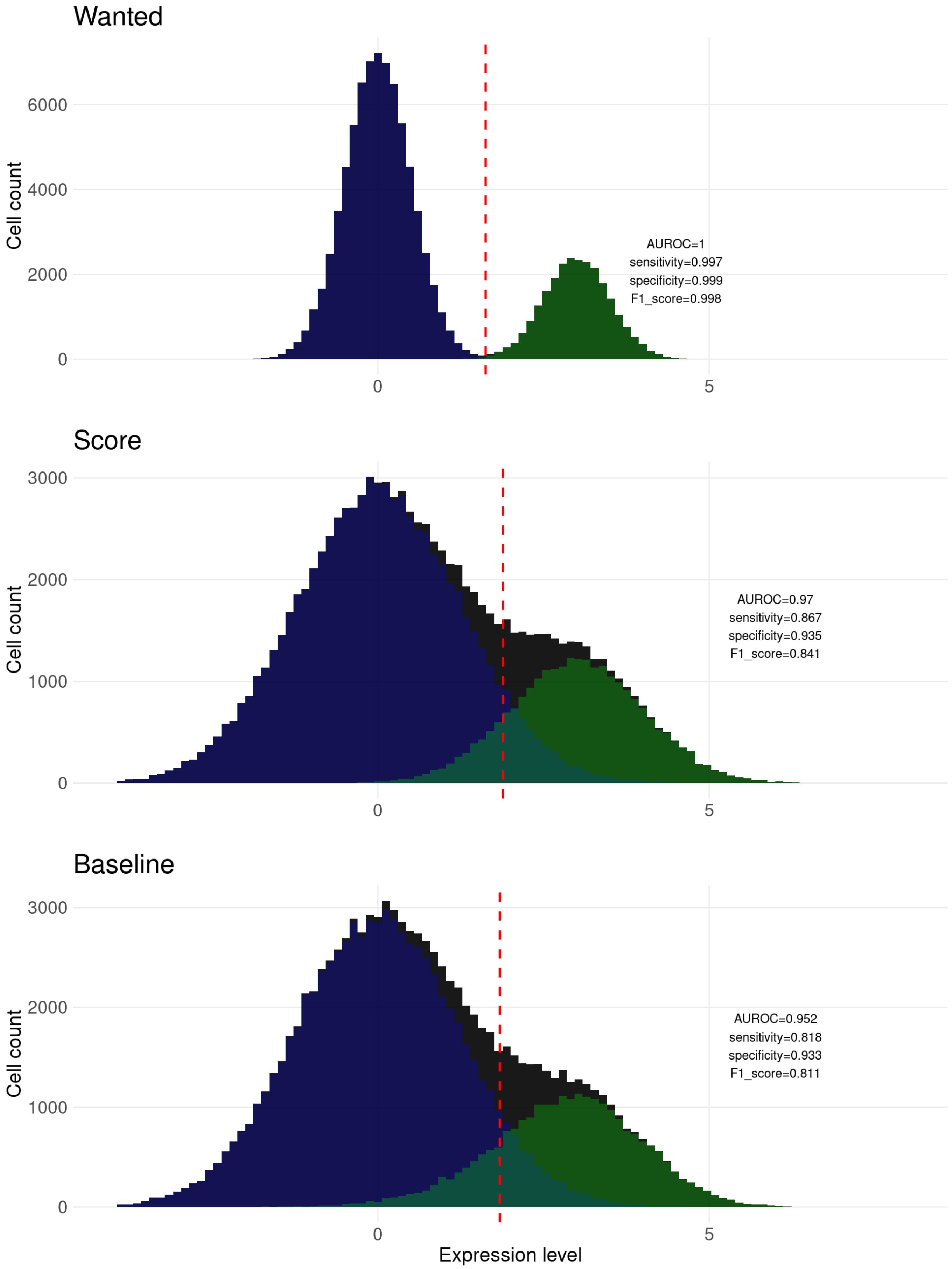


Supplementary Figure 31: Example simulated data showing behaviour of AUROC and F1-score on separation of positive and negative cells for three scenarios: Wanted, Score, and Baseline.
